# *Botrytis cinerea* loss and restoration of virulence during *in vitro* culture follows flux in global DNA methylation

**DOI:** 10.1101/059477

**Authors:** James Breen, Luis Alejandro Jose Mur, Anushen Sivakumaran, Aderemi Akinyemi, Michael James Wilkinson, Carlos Marcelino Rodriguez Lopez

## Abstract

- Pathogenic fungi can lose virulence after protracted periods of culture but little is known of the underlying mechanisms. Here we present the first single-base resolution methylome for the plant pathogen *B. cinerea* and identify differentially methylated genes/genomic regions associated with virulence erosion.
- Cultures were maintained for eight months with subcultures and virulence testing every month. Methylation-sensitive amplified polymorphisms were performed at monthly intervals to characterise global changes to the pathogen’s genome during culture and also on DNA from mycelium inoculated onto *Arabidopsis thaliana* after eight months in culture. Characterisation of culture-induced epialleles was assessed by whole-genome re-sequencing and whole-genome bisulfite sequencing.
- Virulence declined with time in culture and recovered after inoculation on *A. thaliana*. Variation detected by methylation-sensitive amplified polymorphisms followed virulence changes during culture. Whole-genome (bisulfite) sequencing showed marked changes on global and local methylation during culture but no significant genetic changes.
- We imply that virulence is a non-essential plastic character that is at least partly modified by changing levels of DNA methylation during culture. We hypothesise that changing DNA methylation during culture may be responsible for the high virulence/low virulence transition in *B. cinerea* and speculate that this may offer fresh opportunities to control pathogen virulence.

## Introduction

*Botrytis cinerea* is a pathogenic ascomycete responsible for grey mould on a diversity of plant tissue types across hundreds of dicotyledonous plant species (Fournier *et al.*, 2013). It is estimated that this fungus causes annual losses of up to $100 billion worldwide (Weiberg *et al.*, 2013). The wide variety of tissues and species infected by *B. cinerea* suggests it is highly plastic in its ability to penetrate a host thanks to a large ‘arsenal of weapons’ at its disposal. *B. cinerea* is a capable saprotroph and necrotroph, with different genetic types often showing a trade-off between saprotrophic and necrotrophic capabilities (Martinez *et al.*, 2005). As with other pathogens, *B. cinerea* undergoes complex transcriptional and developmental regulation to orchestrate interactions with its host. However, virulence levels of *B. cinerea* strains are not necessarily a fixed feature. For example, virulence has been observed to diminish during protracted *in vitro* culture (Pathirana *et al.*, 2009). Degenerated cultures have been reported in a wide range of pathogenic fungi (Butt *et al.*, 2006), although very little is known about why cultures lose virulence. Several factors have been postulated as possible causes of virulence erosion including dsRNA mycoviruses (Chu *et al.*, 2002; Castro *et al.*, 2003), loss of conditional dispensable chromosomes (Akamatsu *et al.*, 1999; Hatta *et al.*, 2002) or culture-induced selection of nonvirulent strains. However, it is difficult to reconcile mycoviruses infection or chromosome loss as possible causes with a characteristic common to almost all *in vitro*-derived non-virulent fungal strains; that virulence is restored after one passage on their host (Butt *et al.*, 2006). Furthermore, cultured fungal strains lose virulence irrespective of whether the parent culture originated from a single or multiple-spore colony (Butt *et al.*, 2006), discounting the additional possibility of culture-induced selection favouring nonvirulent strains.

The reversibility of this aspect of the observed phenotype comes in response to changes in the growing environment and so should be viewed as a plastic feature of the pathogen (Kelly *et al.*, 2012). In 1942 C.H. Waddington first proposed the term epigenotype to describe the interface between genotype and phenotype. Since then, a large body of research has been carried out to better understand the role of epigenetic regulatory systems in shaping the phenotype of higher organisms in fluctuating environments (Geyer *et al.*, 2011; Tricker *et al.*, 2012). Epigenetic processes operate in a number of ways to alter the phenotype without altering genetic code (Bird, 2007). These include DNA methylation, histone modifications, and mRNA editing and degradation by noncoding RNAs. Such processes are intimately entwined and often work in a synergistic way to achieve changes in phenotype (Rodríguez López & Wilkinson, 2015). DNA methylation, and more specifically cytosine methylation (i.e. the incorporation of a methyl group to carbon 5 of the cytosine pyrimidine ring (5-mC)) is the most studied epigenetic mechanism. It is present across many eukaryotic phyla, including plants, mammals, birds, fish, invertebrates and fungi and provides an important source of epigenetic control for gene expression (Su *et al.*, 2011). In plants and animals, DNA methylation is known to be involved in diverse processes including transposon silencing, X-chromosome inactivation, and imprinting (He *et al.*, 2011). In fungi, overall 5-mC content changes during development in *Phymatotrichu omnivorum* (Jupe *et al.*, 1986) and *Magnaporthe oryzae* (Jeon *et al.*, 2015). Global patterns in DNA methylation has also been shown to dramatically change in lichen fungi species when exposed to the algal symbiont (Armaleo & Miao, 1999). Whole-genome bisulfite sequencing (WGBS) in Ascomycetes has revealed that this group of fungi present repeated loci silenced by methylation but also active genes emthylated within exonic regions. Zemach *et al.* (2010) reported a correlation between gene body methylation and *gene* expression levels in *Uncinocarpus reesii*. More recently, Jeon *et al.* (2015) ascribed a developmental role for DNA methylation in *M. oryzae.* Recent years have seen a dramatic increase in the depth of understanding of how epigenetic control mechanisms operate during plant/pathogen interactions (Boyko & Kovalchuk, 2011; Weiberg *et al.*, 2013). However, little is known of the possible role played by DNA methylation in moderating virulence in fungal plant-pathogen interactions or in its potential role in mediating culture-induced loss of virulence. Here, we explore possible links between erosion of pathogenicity from *B. cinerea* during protracted *in vitro* culture and dynamics of DNA methylation across their genomes. We used Methylation Sensitive Amplified Polymorphisms (MSAPs) (Reyna-López *et al.*, 1997) to provide a preliminary survey of methylome flux associated with gradual reduction in *B. cinerea* pathogenicity with time culture. We next sought to identify Differentially Methylated Regions (DMRs) associated with progressive loss of pathogenicity during *in vitro* culture using WGBS.

## Materials and Methods

### *Botrytis cinerea* culture and inoculation

Seven *Botrytis cinerea* cultures (IMI169558 isolate (Thomma *et al.*, 1999)) were initiated from a single frozen inoculum and cultured and harvested for 32 weeks as described in Johnson *et al.* (2007). After 4 weeks in culture (T0) the initial culture was subcultured to 7 plates containing fresh medium. Mycelium from each plate was subsequently subcultured every 4 weeks (1 month hereafter) into fresh medium. A mycelium sample was taken from all replicates for DNA extraction and conidia harvested from five replicates at every subculture for virulence analysis. Finally, after the T8 challenge, *B. cinerea* was isolated from the infected areas, cultured and immediately used to challenge *A. thaliana* plants to test virulence recovery (Figure S1) (T8P).

### Plant material

*A. thaliana* Col-0 seeds were obtained from the Nottingham Arabidopsis Stock Centre (NASC) and cultivated in Levington Universal compost in trays with 24-compartment inserts (Johnson *et al.*, 2003). Plants were maintained in Conviron (Controlled Environments Ltd) growth rooms at 24°C with a light intensity of 110 μmol m^-2^/s and an 8 h photoperiod for 4 weeks. For ease of treatment, plants were transferred to Polysec growth rooms (Polysec Cold Rooms Ltd, UK; http://www.polysec.co.uk/), maintained under the same conditions.

### DNA isolation

126 *B. cinerea* genomic DNA (gDNA) extractions (comprised of 2 mycelium samples of each of the 7 plates at *in vitro* time point (T1-T8) and from of the last time point culture transplanted onto to *A. thaliana* (T8P)) were performed using the DNeasy 96 Plant Kit (Qiagen, Valencia, CA) according to manufacturer’s instructions. Isolated DNA was diluted in nanopure water to produce working stocks of 10 ng.μl^-1^. DNA from *B. cinerea* inoculated *A. thaliana* was extracted from five leaf samples at each time point using DNeasy Mini Kit (Qiagen, Valencia, CA) as above. DNA samples were diluted to 1 ng.μl^-1^.

### Scoring *B. cinerea* lesion phenotypes

For assessments of infection phenotypes, leaves from five *A. thaliana* Col-0 plants (leaf stage 7/ 8 as defined by Boyes *et al.* (2001), were inoculated on the adaxial surface with 5 μl of spore suspension collected at each subculture time (T0-T8P). Controls were inoculated with PDB. Disease lesions were assessed 3 days post inoculation. A weighted scoring method was used to categorize lesion phenotypes (Lloyd *et al.*, 2011). High virulence symptoms (water-soaking, chlorosis, and spreading necrosis) were conferred a range of positive scores and the resistant symptoms (necrosis limited to inoculation site) were given negative scores (Figure 1A). A weighted score was produced arithmetically from the lesion scores of replicates. Inoculated *A. thaliana* leaves at T0, T8 and T8P were collected 3 days after inoculation for estimation of *in planta* fungal development by quantitative real-time PCR (qPCR) (Figure S1).

**Figure 1:**
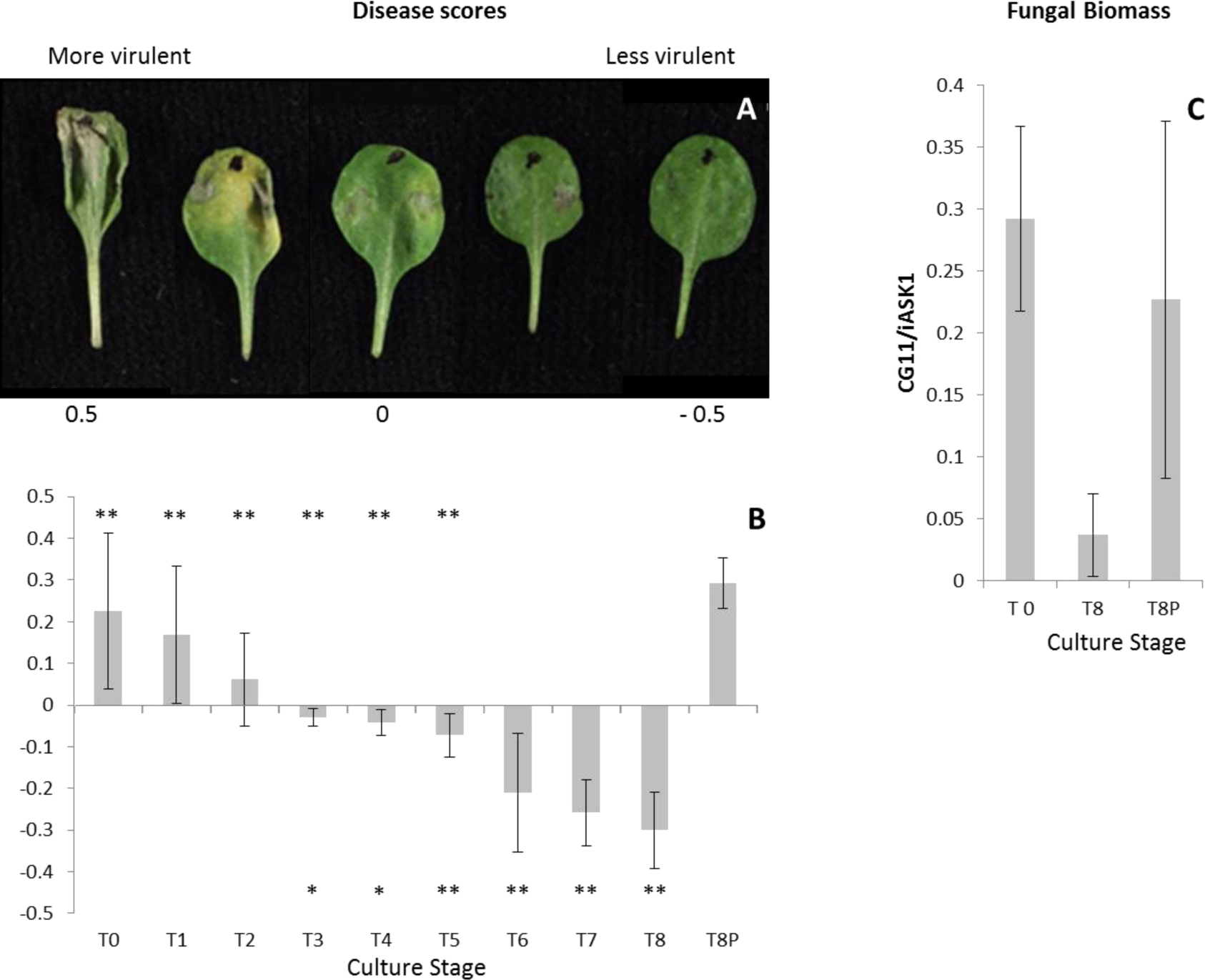
Estimation of *Botrytis cinerea* virulence on *Arabidopsis thaliana*. **(a)** Examples of *A. thaliana* Col-0 leaves drop inoculated with *B. cinerea* and presenting disease scores ranging from −0.5 (low virulence) to 0.5 (high virulence). **(b)** Estimated *B. cinerea* virulence at each time culture point. Virulence of *B. cinerea* cultures was estimated over a period of eight months in culture (T0, initial inoculum; T8, eight months in culture) and after 8 months in culture and a single passage on *A. thaliana* (T8P). A weighted scoring method was used to categorize *B. cinerea* lesion phenotypes 3 days post inoculation. Virulence symptoms (water-soaking, chlorosis, and spreading necrosis) were conferred a range of positive scores and the resistant symptoms (necrosis limited to inoculation site) were given negative scores. Asterisk symbols under the horizontal axis indicate significant differences (*(T-Test; P<0.05) and ** (T-Test; P<0.01)) between T0 and the time point over the asterisk. Asterisk symbols over the horizontal axis indicate significant differences (** (T-Test; P<0.01)) between T8P and the time point under the asterisk. **(c)** Detection of *in planta B. cinerea* hyphal mass in *A. thaliana* Col-0 by qPCR as described by Gachon and Saindrenan, 2004.

### Estimation of *in planta* fungal growth by qPCR

Reaction mixtures for qPCRs (25 μl) were prepared by mixing 10 ng of DNA with 12.5 μl of SYBR™ Green Mastermix (Applied Biosystems, UK) and primers (300 nM final concentration). Arabidopsis and Botrytis primers (Table S1) generated a 131 bp amplicon of the Shaggy-kinase-like gene and a 58 bp amplicon of the Cutinase A gene respectively (Gachon & Saindrenan, 2004). qPCRs were carried out using a Bio-Rad ABI7300 thermocycler using the following conditions: 15 min (95°C) followed by 50 cycles of 15 s (95°C), 30 s (58°C) and 1 min (72°C). This was followed by a dissociation (melting curve), according to the software procedure. Serial dilutions of pure genomic DNA from each species were used to trace a calibration curve, which was used to quantify plant and fungal DNA in each sample. Results were expressed as the CG11/iASK ratio of mock-inoculated samples.

### MSAP procedure

We used MSAPs (Rodríguez López *et al.*, 2012) to reveal global patterns of variability that reflect the divergence in methylation and sequence mutations between *B. cinerea* samples. 14 samples per time point (T2 to T8P) were analysed (2 replicated DNA extractions per culture plate). For each individual sample, 50ng of DNA were digested and ligated for 2 h at 37°C using *Eco*RI (5U) and *Msp*I or *Hpa*II (1U) (New England Biolabs), 0.45 μM *Eco*RI adaptor, 4.5 μM *Hpa*II adaptor (Table S1 for all oligonucleotide sequences) and T4 DNA ligase (1U) (Sigma) in 11 μl total volume of 1X T4 DNA ligase buffer (Sigma), 1μl of 0.5M NaCl, supplemented with 0.5 μl at 1mg/ml of BSA. Enzymes were then inactivated by heating to 75°C for 15 min. Restriction/ligation was followed by two successive rounds of PCR amplification. For preselective amplification, 0.3 μl of the restriction/ligation products described above were incubated in 12.5 μl volumes containing 1X Biomix (Bioline, London, UK) with 0.05 μl of Preamp EcoRI primer and 0.25 μl Preamp HpaII/MspI (both primers at 10 uM) supplemented with 0.1 μl at 1mg/ml of BSA. PCR conditions were 2 min at 72°C followed by 30 cycles of 94°C for 30 s, 56°C for 30 s and 72°C for 2 min with a final extension step of 10 min at 72°C. Selective PCRs were performed using 0.3 μl of preselective PCR product and the same reagents as the preselective amplification but using FAM labelled selective primers (E2/H1). Cycling conditions for selective PCR were as follows: 2 min (94°C), 13 cycles of 30 s (94°C), 30 s (65°C, decreasing by 0.7°C each cycle), and 2 min (72°C), followed by 24 cycles of 30 s (94°C), 30 s (56°C), and 2 min (72°C), ending with 10 min (72°C). Fluorescently labelled MSAP products were diluted 1:10 in nanopure sterile water and 1 μl was combined with 1 μl of ROX/HiDi mix (50 μl ROX plus 1 ml of HiDiformamide, Applied Biosystems, USA). Samples were heat-denatured at 95°C for 3–5 min and snap-cooled on ice for 2 min. Samples were fractionated on an ABI PRISM 3100 at 3 kV for 22 s and at 15 kV for 45 min.

### Analysis of genetic/epigenetic variability during time in culture using MSAP

MSAP profiles were visualized using GeneMapper Software v4 (Applied Biosystems, Foster City, CA). A qualitative analysis was performed in which loci were scored as ‘‘present’’ (1) or ‘‘absent’’ (0) to create a presence/absence binary matrix. MSAP selected loci were limited to amplicons in the size range 80-585bp to reduce potential for size homoplasy (Caballero *et al.*, 2008). Samples were grouped according to the cumulative period in culture at the time of collection Samples collected after 2, 3, 4, 5, 6, 7 and 8 months were denoted T2, T3, T4, T5, T6, T7, T8 and T8P respectively whereas those cultured for 8 months and then inoculated onto onto *A. thaliana* were labelled T8P.

Similarity between MSAP profiles obtained from primer combination E2/H1 and both enzymes (*Hpa*II and *Msp*I) was first visualized using Principal Coordinate Analysis (PCoA) (Gower, 1966) using GenAlex (v.6.4) (Peakall & Smouse, 2012). We then used Analysis of Molecular Variance (AMOVA) (Excoffier *et al.*, 1992) to evaluate the structure and degree of diversity induced by different times in culture. Pairwise PhiPT (Michalakis & Excoffier, 1996) comparisons between samples restricted with *Hpa*II or *Msp*I from each time point and the samples after the first passage (2 months in culture, T2) were used to infer their overall level of divergence with time in culture (i.e., the lower the PhiPT value between samples restricted using *Hpa*II or *Msp*I, the smaller the differentiation induced by culture and the same samples). AMOVA was subsequently calculated using GenAlex (v.6.5) to test the significance of PhiPT between populations (Michalakis & Excoffier, 1996), with the probability of non-differentiation (PhiPT=0) being estimated over 9,999 random permutations.

Mantle test analysis was used to estimate the correlation between the calculated pairwise molecular distances and the difference in virulence between culture time points. The level of significance was estimated over 9,999 random permutations tests, as implemented in Genalex v6.5.

### Methylation analysis by WGBS

DNA from 9 biological replicates from culture of two culture ages (1, 8 months) and from 9 replicates of T8P were randomly selected for sequencing. Biological replicates were used to generate 3 pooled samples per culture age. Bisulphite treatment was performed independently from 50ng of genomic DNA of each pooled sample using the EZ DNA methylation-Gold™ Kit (Zymo Research) according to the manufacturers’ instructions but adjusting the final column purification elution volume to 10 μl. Following Bisulphite treatment, recovered DNA from each pool was used to estimate yield using a NanoDrop 100 spectrophotometer using the RNA setting. Bisulphite treated samples were then used to create a sequencing library using the EpiGnome™ Methyl-Seq Kit and Index PCR Primers (4-12) according to manufacturer’s instructions. In order to provide a reference draft sequence for the alignment of the bisulphite treated DNA and to detect any culture induced genetic mutations, 10 ng of native (non-bisulphite treated) DNA extracted from cultures from time points 1 month (T1) and 8 month (T8) were sequenced and compared to the reference B. *cinerea* B05.10 genome sequence. Libraries were prepared using the EpiGnome™ Methyl-Seq Kit and Index PCR Primers (Epicentre) (1-2) according to manufacturer’s instructions.

Library yield was determined by Qubit dsDNA High Sensitivity Assay Kit. Agilent 2100 Bioanalyzer High-Sensitivity DNA Chip was used to assess library quality and determine average insert size. Libraries were then pooled and sequenced on Illumina HiSeq2000 (Illumina Inc., San Diego, CA) 100bp paired-end V3 chemistry. Data available at Sequence Read Archive (PRJEB14930).

### Sequence analysis and differential methylation analysis

Sequencing reads were trimmed to remove adaptors using *TrimGalore*! (<u>http://www.bioinformatics.babraham.ac.uk/projects/trim_galore</u>) and *Cutadapt* (Martin, 2011). Whole genome re-sequencing reads were aligned to the published genome from *B. cinerea* B05.10 (Broad Institute’s *B. cinerea* Sequencing Project) using *bowtie2* (Langmead & Salzberg, 2012) and variants were called using *freebayes* (Garrison & Marth, 2012) after filtering for coverage greater than 10x and >30 variant quality and removal of multi-allelic variants, variants with missing data in one sample and variants with an observed allele frequency less than 0.5 (Jeon *et al.*, 2013). Variant categories were analysed using *CooVar* (Vergara *et al.*, 2012), *bedtools* (Quinlan & Hall, 2010) and custom scripts. Bisulfite treated libraries were mapped using Bismark (Krueger and Andrews, 2011) and bowtie2, duplicates removed and methylation calls extracted using samtools (Li *et al.*, 2009) and in-house scripts (available upon resquest). Bisulfite sequencing efficiency was calculated by aligning reads to the B. *cinerea* mitochondrial genome scaffold (B05.10) and identifying non-bisulfite converted bases. Differentially methylated regions were called using a sliding window approach described in swDMR (<u>https://code.google.com/p/swdmr/</u>). The significance of the observed DMRs was determined using a three sample Kruskal-Wallis (3 sample) test between T1, T8 and T8P.

## Results

### Pathogenicity analysis of *Botrytis cinerea*

Isolates from all culture time points produced lesions on *A. thaliana* leaves but they varied in their severity (Figure 1A). There was a progressive decline in disease scores recorded over the eight-month period (i.e. T0-T8) (Figure 1B), with loss of virulence becoming significant from 3 months (T3) onwards (T-Test P<0.05) (Figure 1B). The disease scores for the T8P challenge did not differ significantly from those recorded at T0 culture time, indicating that virulence had recovered following a single passage through a plant (Figure 1B). Conversely, T8 virulence scores were significantly lower than those obtained from T0 to T5 and also than T8P (T-Test P<0.05) (Figure 1B). Infected leaves with T0 and T8P cultures did not show significant differences in fungal DNA content measured by qPCR (Figure 1C). However, both showed significantly higher levels of fungal DNA (T-Test P<0.05) than those infected using T8 cultures (Figure 1C).

### Analysis of genetic and epigenetic variance during culture using MSAPs

MSAP profiles generated 74 scored loci across the 112 samples of eight *B. cinerea* culture times used in this study (T2-T8P). Multivariate analysis revealed that the MSAP profiles of *B. cinerea* became progressively more dissimilar to the first time point analysed (T2) with increasing culture age (Figure 2). Both, PCoA (Figure 2A and C) and PhiPT values (Figure 2B) showed higher levels of culture-induced variability when using *Hpa*II than when using *Msp*I. PCoA shows that samples cultivated for 3, 4, 5, 6, and 7 months occupied intermediate Eigenspace between samples cultivated for 2 and 8 months (Figure 2C). Furthermore, a partial recovery of the MSAP profile was observed on samples cultured for 8 months after one fungal generation on the host plant (T8P) (Figure 2C). Calculated PhiPT values between each time point and T2 samples show a progressive increase in MSAP profile distance with time in culture when samples were restricted with both enzymes. Analysis of Molecular Variance (AMOVA) analysis shows that the calculated PhiPT values were significantly different (P<0.05) between T2 and time points T6, T7, T8 and T8P when using *Msp*I and T7, T8 and T8P when using *Hpa*II (Figure 2B). Mantle test analysis showed significant correlations between disease score differences among culture times and pairwise PhiPT values from MSAP profiles generated using *Msp*I (R^2^ = 0.316; P=0.005) and *Hpa*II (R^2^ = 0.462; P=0.002) (Figure S2).

**Figure 2:**
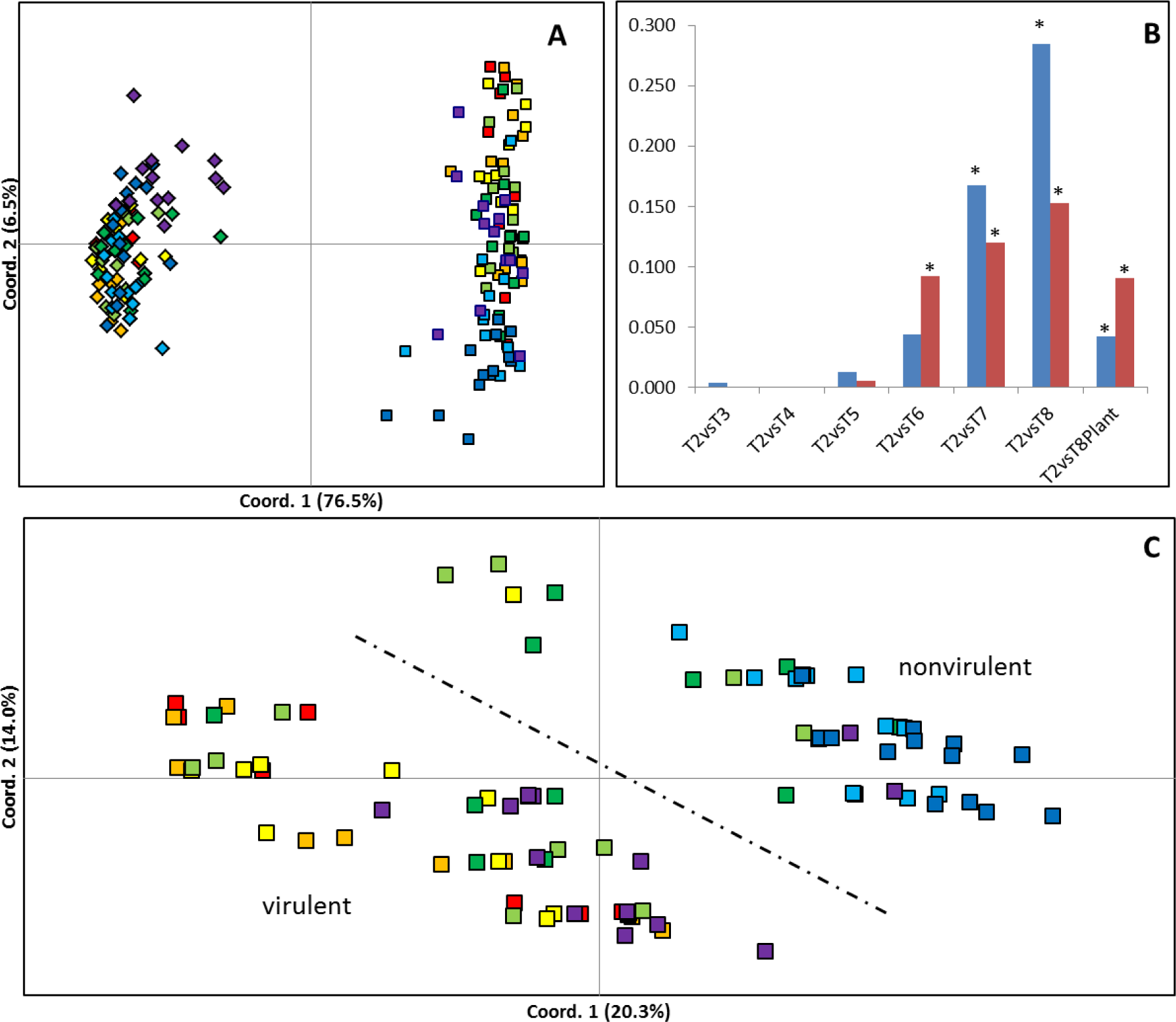
Effect of time in culture on genetic/epigenetic instability. **(a, c)** Principal coordinate diagrams based on the Euclidian analysis of methylation-sensitive amplified polymorphisms (MSAP) using enzymes *Hpa*II (squares) and *Msp*I (romboids) **(a)** and using enzyme *Hpa*II distances **(c)**. 14 replicates from each time point are represented as red (T2: 2 months in culture), orange (T3: 3 months in culture), yellow (T4: 4 months in culture), light green (T5: 5 months in culture), dark green (T6: 6 months in culture), light blue (T7: 7 months in culture), dark blue (T8: 8 months in culture), and purple (T8P: 8 months+plant). The dashed line separates samples with higher average levels of virulence from those of lower average levels of virulence. **(b)** Calculated Pairwise PhiPT (Michalakis & Excoffier, 1996) comparisons between samples restricted with *Hpa*II (Blue) or *Msp*I (Red) from each time point and the samples after the second passage (2 months in culture). * Indicates significantly different PhiPT values between T2 and the time point under the asterix based on 10,000 permutations (P = 0.05).

### *B. cinerea* genome resequencing

Divergence in MSAP profiles may occur through changes in methylation or through genetic mutation. However, the recovery of profile affinity towards that of the original inoculum after a single passage on Arabidopsis is difficult to reconcile genetically, suggesting the majority of the changes to MSAP profiles was caused by differences in DNA methylation rasther than sequence mutations. We therefore sought to better characterise mutational changes using a genome-wide sequencing approach. DNA extractions from two time points (1 month (T1) and 8 months (T8) in culture) were compared to the Broad Institute’s *B. cinerea* B05.10 reference genome sequence and used to assess mutational change during culture. Both samples were sequenced to an average depth of 37.47x (35.64x for the 1-month culture and 39.30x for the 8-month culture) with an average of 80% of reference bases being sequenced to a depth greater than 10x. After filtering we found 2,331 sequence variants after 8 months in culture, of which 1,030 (44%) were small insertions and deletions (INDELs) and 1,301 (56%) SNPs. Of these, just 454 were located within genes including: 251 synonymous variants, 198 non-synonymous mutations (193 causing missense variations and 5 causing premature stop codons (non-sense mutations). An additional 5 variants that altered predicted splice junctions were also identified.

We next focused the search for variants within the sequence of 1,577 *B. cinerea* genes with known function including: secondary metabolism, conidiation, sclerotium formation, mating and fruit body development, apoptosis, housekeeping, signalling pathways (Amselem *et al.*, 2011) and virulence *senso lato* genes. The virulence *sensu lato* genes included: appressorium-associated genes (Amselem *et al.*, 2011), virulence *sensu stricto* genes (Choquer *et al.*, 2007) and plant cell wall disassembly genes (CAZyme genes) (Blanco-Ulate *et al.*, 2014). We found 68/1,577 (4.3%) of the tested genes contained one or more variants between T1 and T8 (See Table 1 and Table S2 for a comprehensive list of genes with variants).

**Table 1:**
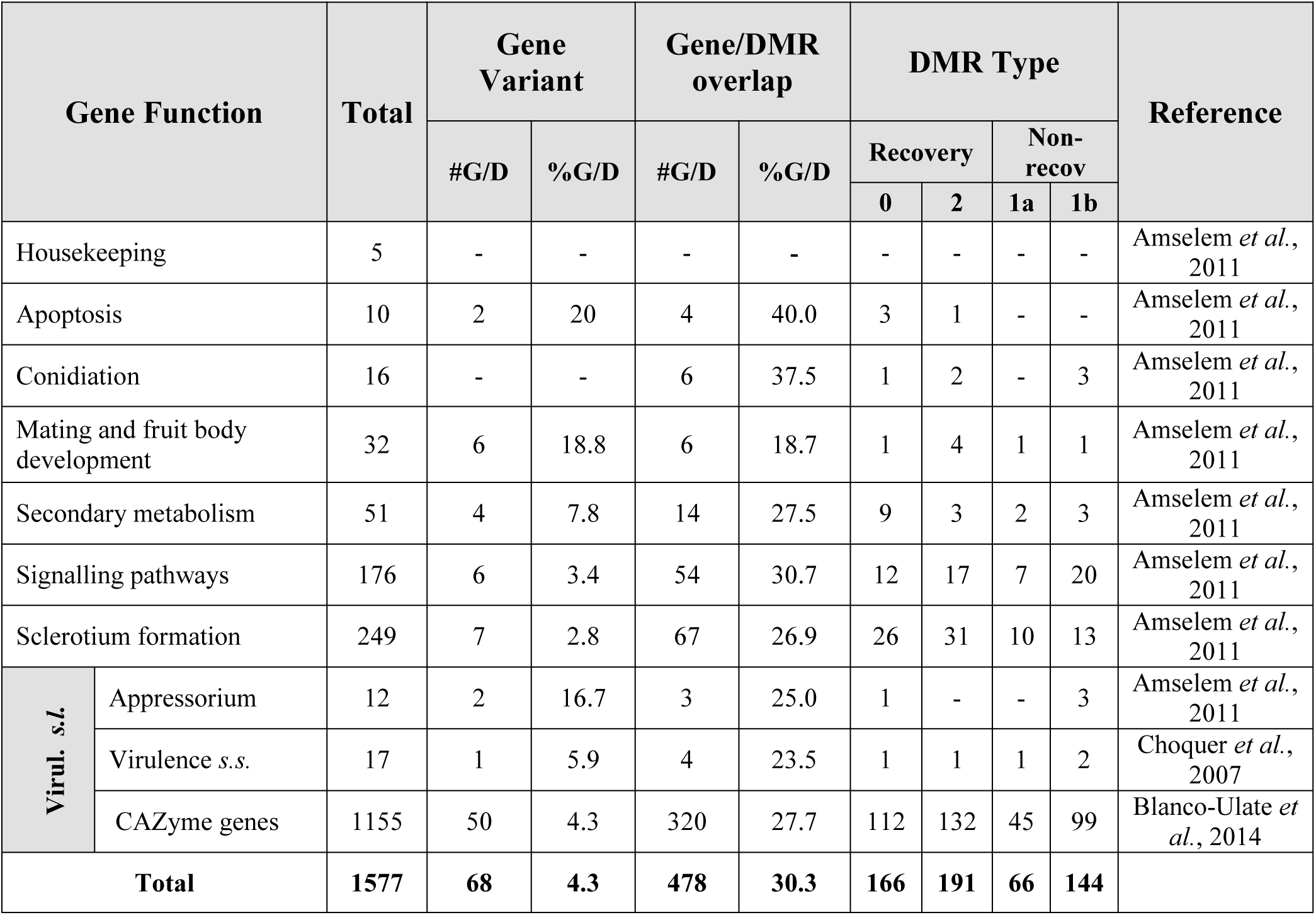
*Botrytis cinerea* virulence genes with known function overlapping with *in vitro* culture induced DMRs. **Columns Gene Variant** and Gene/DMR overlap indicate the number (#) and percentage (%) of genes in each functional group with a genetic variant or overlapping with a Differentially Methylated Region respectively. Column DMR type indicate how methylation levels changed when comparing all three samples (T1, T8 and T8P). DMRs were grouped according to their changing patterns into recovery (T1=T8P) and non-recovery (T1< >T8P). Two subgroups where found for recovery (T1=T8P<T8 (Type 0) and T1=T8P > T8 (Type 2)) and non-recovery (T1>T8=T8P (Type 1a) and T1=T8<T8P (Type 1b)). Virul. s.l. indicates virulence genes in abroad sense and include genes associated to: Appressorium formation, virulence on a strict sense and CAZyme genes.

| Gene Function |  | Total | Gene Variant |  | Gene/DMR overlap |  | DMR Type |  |  |  | Reference |
| --- | --- | --- | --- | --- | --- | --- | --- | --- | --- | --- | --- |
|  |  |  | #G/D | %G/D | #G/D | %G/D | Recovery |  | Non-recov |  |  |
|  |  |  |  |  |  |  | 0 | 2 | 1a | 1b |  |
| Housekeeping |  | 5 | - | - | - | - | - | - | - | - | Amselem *et al.*, 2011 |
| Apoptosis |  | 10 | 2 | 20 | 4 | 40.0 | 3 | 1 | - | - | Amselem *et al.*, 2011 |
| Conidiation |  | 16 | - | - | 6 | 37.5 | 1 | 2 | - | 3 | Amselem *et al.*, 2011 |
| Mating and fruit body development |  | 32 | 6 | 18.8 | 6 | 18.7 | 1 | 4 | 1 | 1 | Amselem *et al.*, 2011 |
| Secondary metabolism |  | 51 | 4 | 7.8 | 14 | 27.5 | 9 | 3 | 2 | 3 | Amselem *et al.*, 2011 |
| Signalling pathways |  | 176 | 6 | 3.4 | 54 | 30.7 | 12 | 17 | 7 | 20 | Amselem *et al.*, 2011 |
| Sclerotium formation |  | 249 | 7 | 2.8 | 67 | 26.9 | 26 | 31 | 10 | 13 | Amselem *et al.*, 2011 |
| Virul. s.l. | Appressorium | 12 | 2 | 16.7 | 3 | 25.0 | 1 | - | - | 3 | Amselem *et al.*, 2011 |
|  | Virulence s.s. | 17 | 1 | 5.9 | 4 | 23.5 | 1 | 1 | 1 | 2 | Choquer *et al.*, 2007 |
|  | CAZyme genes | 1155 | 50 | 4.3 | 320 | 27.7 | 112 | 132 | 45 | 99 | Blanco-Ulate *et al.*, 2014 |
| Total |  | 1577 | 68 | 4.3 | 478 | 30.3 | 166 | 191 | 66 | 144 |  |

### Characterisation of DNA Methylation changes by WGBS

We next characterised genome-wide DNA methylation flux by conducting WGBS from triplicated genomic DNA extractions from mycelia of two different culture ages (T1 and T8) and eight month cultures after inoculation onto *A. thaliana* plant (T8P). This yielded 187.5 million reads, ranging from 12.61 to 34.05Gbp of sequence per sample after quality filtering. Mapping efficiency of each replicate ranged from 54.3 to 67.6%, resulting in samples that generated between 33 and 55x coverage of the 42.66Mbp genome (Table S3). This is the highest genome coverage achieved to date for any fungal species after bisulfite sequencing. For each sample, we covered over 91-93% of all cytosines in the genome (Table S3), with all samples having at least 81% of cytosines covered by at least four sequencing reads, allowing methylation level of individual sites to be estimated with reasonable confidence.

WGBS identified an average of 15,716,603 mC per sample, indicating an average methylation level of 0.6% across the genome as a whole (Table S4). The most heavily methylated context was CHH followed by CG and CHG (where H is A, C or T) (Table S4). Global levels of mC did not significantly change with culture time, although methylation in the rarer CG and CHG contexts increased significantly (T-test, p=0.0008 and 0.0018) between 1 and 8 months in culture (Figure 3A). However, modest declines were apparent in both contexts on eight month old cultures inoculated onto *A. thaliana* (T8P) (Figure 3A) they proved not significant ((T-test, p>0.05). Analysis of local levels of DNA methylation across the largest *B. cinerea* contig (Supercontig 1.1) showed that DNA methylation is unevenly distributed and locally clustered (Figure 3B-D). Observed clustering patterns were similar in all analysed samples (Figure 3B-D).

**Figure 3.**
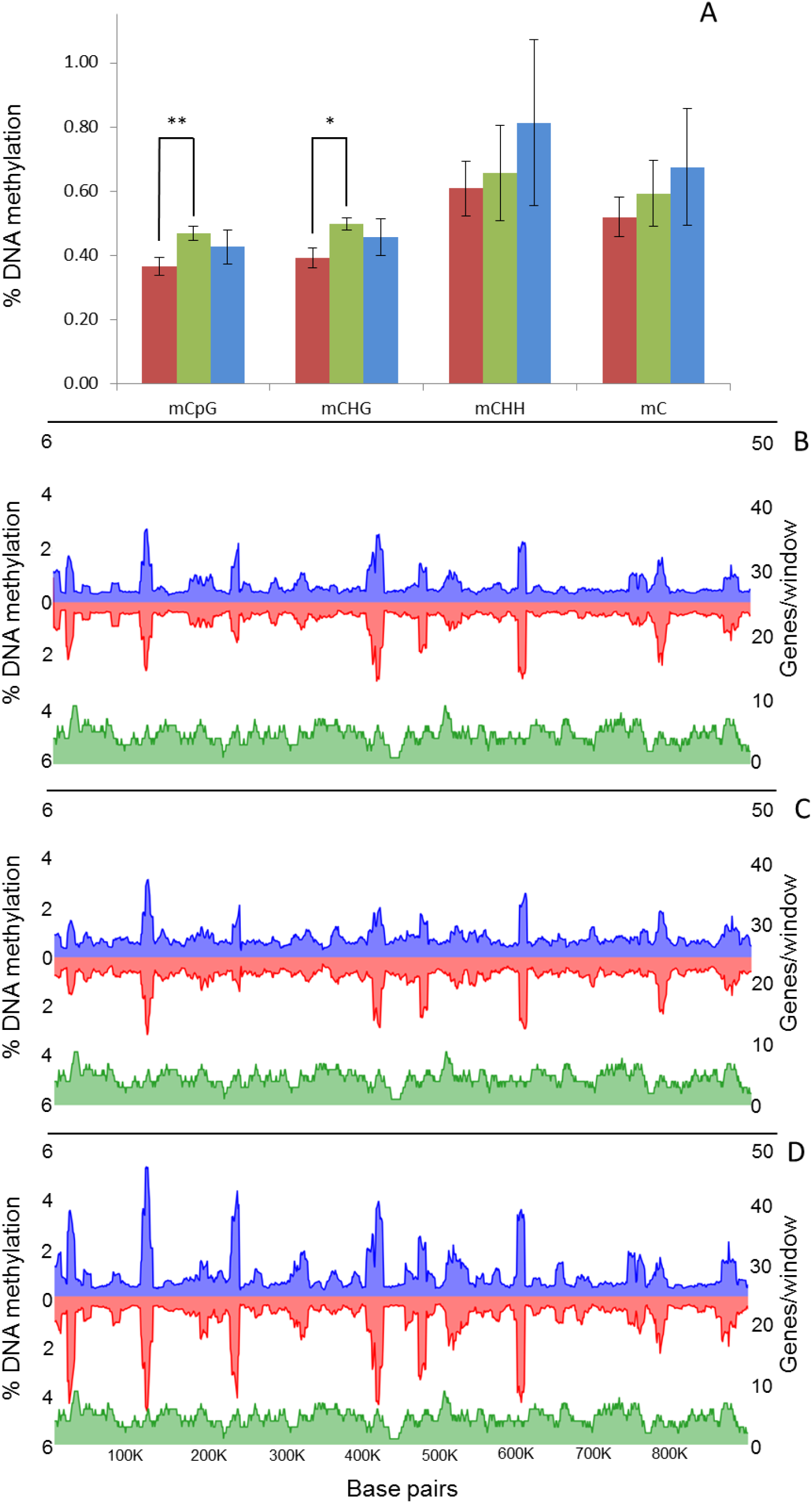
Global changes on genomic distribution and levels of DNA methylation in B. cinerea. **A)** Global average percentage of mCs ((number of mCs/total Cs)*100)) at each time point (T1, red; T8, green and T8P, blue). **B-D)** Methylcytosines (mCs) density from each strand (blue, positive and red, negative strand) across supercontig 1.1 at each time point (**B**=T1; **C**=T8 and **D**=T8P) was calculated and plotted as the percentage of mCs (as above) in each 10kb window.

We next investigated the fine-scale distribution of DNA methylation across genic and regulatory regions by surveying the location and density of mCs on all B. cinerea genes (exons and introns), and presumed promoter regions, defined here as 1.5 kb upstream of the Transcription Starting Site (TSS). Among these, mC levels increased between 1,000 and 500bp upstream of TSS. This was followed by a sharp decrease then and increase in methylation before and after the start of the coding sequence respectively (Figure 4). When time points T1, T8 and T8P were compared on this genomic context, methylation levels where higher on T8P samples (Figure 4), in parallel with that observed at a whole-genome level (Figure 3A).

**Figure 4:**
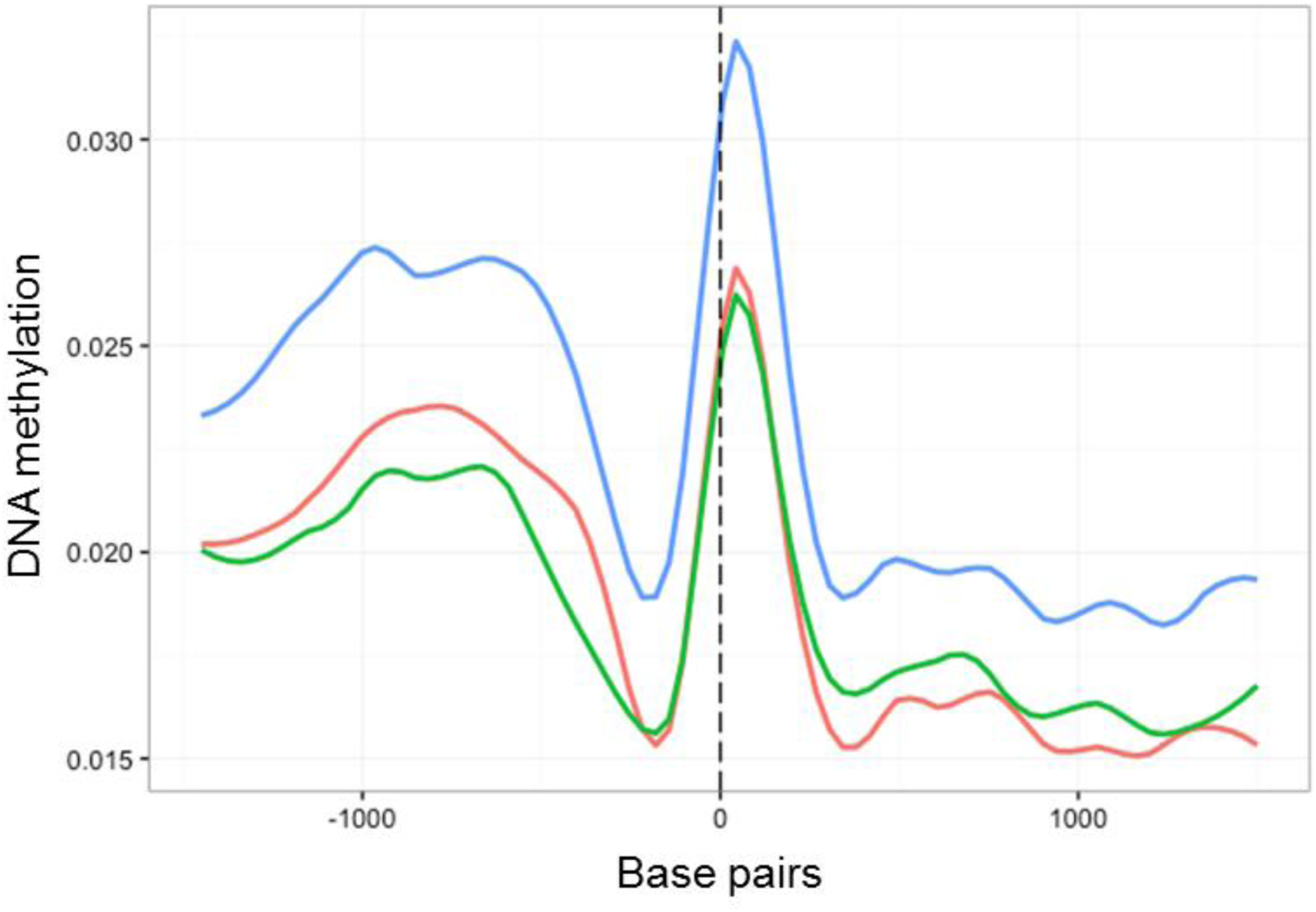
Methylation density across transcription start sites (TSS) regions of all *B. cinerea* selected genes. Methylation level (Vertical axis) at each time point (T1, red; T8, green and T8P, blue) was identified as the proportion of methylated cytosines against all cytosines in 30bp windows 1.5kb before and after the transcription start site.

The same methylation density analysis was then carried out for five housekeeping genes in Botrytis (i.e., G3PDH (BC1G_09523.1); HSP60 (BC1G_09341.1); Actin (BC1G_08198.1 and BC1G_01381.1) and Beta Tubulin (BC1G_00122.1). All five loci possessed low levels of DNA methylation in every context and there were no changes in DNA methylation between culture time points (i.e., T1, T8 and T8P) (Data not shown).

In common with most genes, those encoding putative CAZymes, proteins secreted by *B. cinerea* upon plant infection (Table S5) (Blanco-Ulate *et al.*, 2014), contained more methylation in regions immediately upstream of the TSS (Figure 5A). However, these genes showed higher levels of methylation after 8 months in culture than at T1 and at T8P. The observed increase in methylation on T8 samples was primarily driven by increased methylation in the CHG and CG (Figure 5B-C) contexts. In contrast, CHH sites (Figure 5D) showed highst levels of methylation on the T8P samples; following the general trend observed both at a whole genome level (Figure 3A) and by all *B. cinerea* genes (Figure 4).

**Figure 5:**
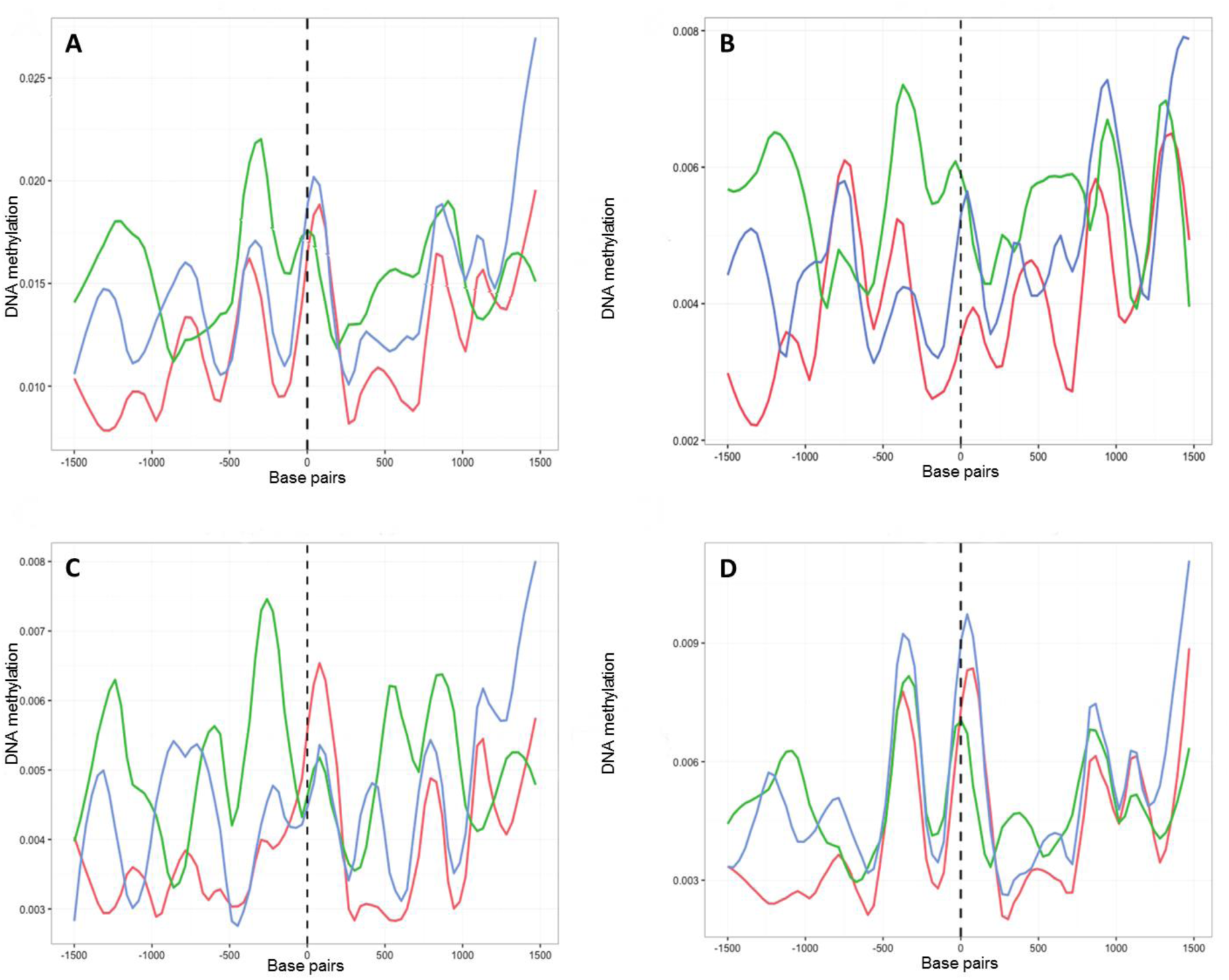
Methylation density across transcription start sites (TSS) found in 131 genes associated to polysaccharide degradation in *B. cinerea* (27). Methylation level (Vertical axis) at each time point (T1, red; T8, green and T8P, blue) was identified as the proportion of methylated cytosines against all cytosines in 30bp windows 1.5kb before and after the transcription start site (TSS) (Horizontal axis). **A)** Methylation level for all mCs; **B)** Methylation level for CpG context; **C)** Methylation level for CpHpG context and **D)** Methylation level for CpHpH context.

### Detection of culture induced DMRs

We sought to identify DMRs between three culture times (T1, T8 and T8P) by comparing methylation levels across the whole genome of all samples implementing swDMR sliding window analysis. Analysis of DMR length distribution showed DMRs sizes ranging from 12 to 4994bp (Figure 6A). The sliding window approach identified 2,822 regions as being significantly differentially methylated in one of the samples compared to the other two for all mCs (Table 2, Table S6). Overall methylation levels of DMRs decreased in all contexts (CG, CHG and CHH) as time in culture progressed (from T1 to T8) but this was followed by a recovery of DNA methylation levels after 8 month cultures were inoculated onto *A. thaliana* (T8P) (Figure 6b). However, it is worth noting that T8 showed a larger number of outlier DMRs (greater than two standard deviations away from the mean) that exhibited significantly higher levels of methylation than the average (Figure 6B).

**Figure 6:**
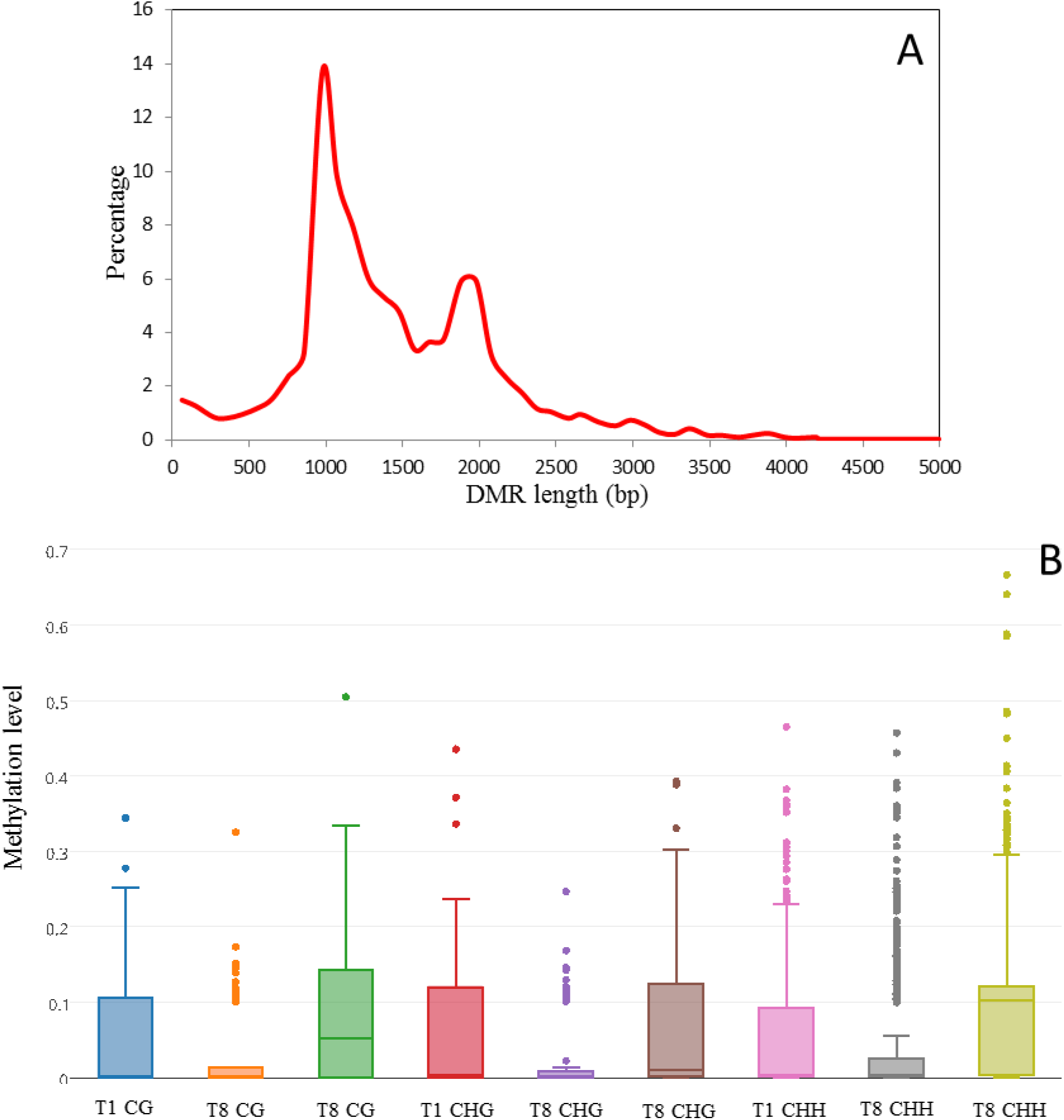
Analysis of *in vitro* culture induced DMRs in *Botrytis cinerea*. **(a)** Length distribution of *in vitro* culture induced DMRs (i.e., regions presenting significantly different methylation levels between one sample and the other two samples (FDR<0.01)) in *Botrytis cinerea.* DRMs were determined using a three sample Kruskal-Wallis test. Methylation levels were analysed by sliding window analysis using swDMR. **(b)** Methylation level distribution in *in vitro* induced DMRs. Boxplot of 3 sample sliding window differential methylation analysis using swDMR. The boxplots shows the distribution of each methylation in each context (CG, CHG and CHH) at three time points (T1, T8 and T8P), circles indicate DMRs with outlier levels of methylation.

**Table 2:**
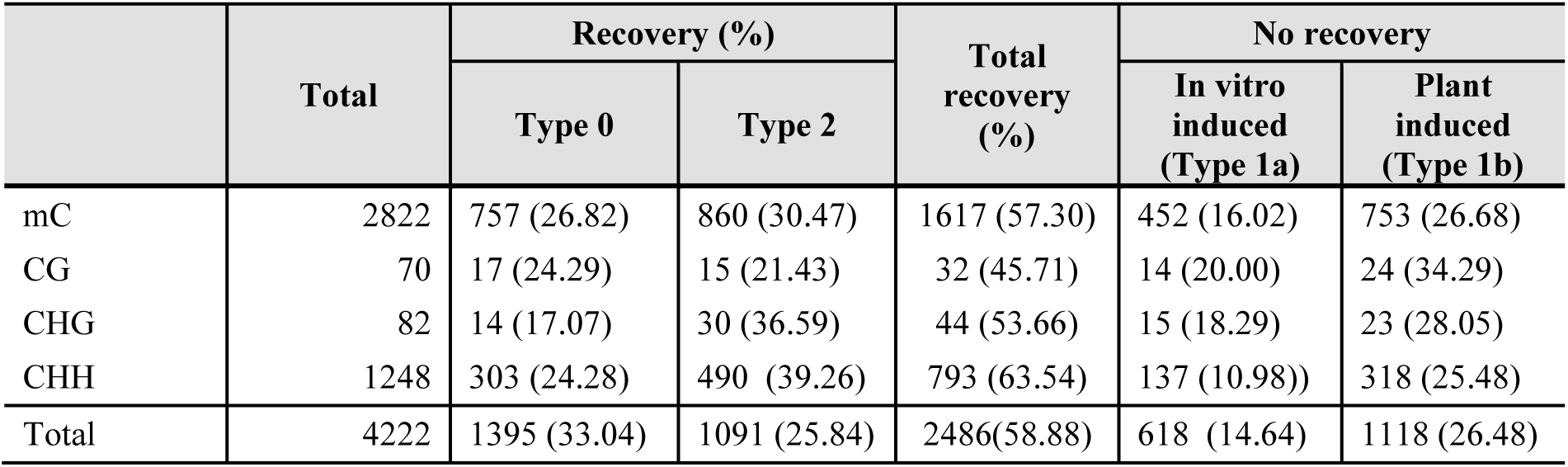
Number of Differentially Methylated Regions (DMRs) between T1, T8 and T8P samples. DRMs (i.e., regions presenting significantly different methylation levels between one sample and the other two samples (FDR<0.01)) were determined using a three sample Kruskal-Wallis test. Methylation levels were analysed by sliding window analysis using swDMR. DMRs were determined for all cytosines and for three methylation contexts. DMRs were grouped according to their changing patterns into recovery (T1=T8P) and non-recovery (T1< > T8P). Two subgroups where found for recovery (T1=T8P < T8 (Type 0) and T1=T8P > T8 (Type 2)) and non-recovery (T1>T8=T8P (Type 1a) and T1=T8<T8P (Type 1b)). Percentage of the total DMRs for each pattern type/sequence context is shown in parenthesis.

When examined individually, variance in DMRs between samples comprised of two main pattern types (Table 2, Table S6): **1.** 57.3% of the detected DMRs showed a recovery pattern inoculation on *A. thaliana* such that there was no difference between T1 and T8P samples but methylation levels diverged significantly in T8 (T1=T8P< >T8 (FDR<0.01))). **2.** The remaining DMRs showed a non-recovery pattern (i.e., T1< >T8P (FDR<0.01)). Two subtypes where found for DMRs showing a DNA methylation recovery pattern: **1.** DMRs showing an increase in methylation with time in culture (T1=T8P<T8 (26.82%) (Type 0 hereafter) and **2.** DMRs showing a decrease in methylation level with time in culture (T1=T8P>T8 (30.47%)) (Type 2). Equally, non-recovery DMRs can be divided into two categories: **1.** DMRs showing a decrease in methylation level with time in culture and no change in methylation level following inoculation on *A. thaliana* (T1>T8=T7P (16.02%)) (Type 1a) and **2.** DMRs not showing changes in methylation level during culture but an increase in methylation after inoculation on *A. thaliana* (T1=T8<T8P (26.68%)) (Type 1b). Curiously, no DMRs were observed to show a progressive increase in methylation level with time in culture and no change in methylation level after inoculation on *A. thaliana* (T1<T8=T8P).

The vast majority (84.5%) of detected DMRs overlapped with 3,055 genic regions in the *B. cinerea* genome (Table 3) while 438 (15.5%) mapped to intergenic regions. Almost all of the genes implicated (98%) included DMRs within the gene body itself, with a small majority (53.9%) also overlapping with the promoter region (Table 3, Table S7). The same analyses were carried out to detect DMRs for CG, CHG and CHH contexts, identifying 70, 82 and 1,248 DMRs respectively for each context (Table 3, Table S7). Of these, 91.4% (CG), 89.0% (CHG) and 85.2% (CHH) overlapped with 68, 84 and 1,339 genes respectively (Table 3, Table S7).

**Table 3:** *Botrytis cinerea in vitro* induced Differentially Methylated Regions overlapping with genes. DRMs overlapping with genes (i.e., regions presenting significantly different methylation levels between one sample and the other two samples (FDR<0.01)) were determined using a three sample Kruskal-Wallis test. Methylation levels were analyzed by sliding window analysis using swDMR. DMRs were determined for all cytosines and for three methylation contexts. DMRs were grouped according to the genic region they overlapped with (i.e, Promoter, promoter and Gene body, promoter, Gene body and 3’UTR, gene body and 3’UTR and gene body). (**) Percentage of the total DMRs overlapping with each particular genic region. (*) Percentage of the total number of genes showing a methylation recovery pattern.

|  | Total genes | Promoter,<br>Gene body<br>and 3'UTR<br>(*) | Promoter<br>only (*) | Promoter<br>and Gene<br>body (*) | Gene body<br>and 3'UTR<br>(*) | Gene body<br>(*) |
| --- | --- | --- | --- | --- | --- | --- |
| mC | 3055 | 626 (20.49) | 61 (2.00) | 1022 (33.45) | 923 (30.21) | 423 (13.85) |
| CG | 68 | 3 (4.41) | 0 (0.00) | 24 (35.29) | 21 (30.88) | 20 (29.41) |
| CHG | 84 | 8 (9.52) | 0 (0.00) | 29 (34.52) | 27 (32.14) | 20 (23.81) |
| CHH | 1339 | 248 (18.52) | 32 (2.39) | 443 (33.08) | 413 (30.84) | 203 (15.16) |
|  | Recovery<br>genes (**) |  |  |  |  |  |
| mC | 1713 (56.07) | 345 (20.14) | 43 (2.51) | 545 (31.82) | 537 (31.35) | 243 (14.19) |
| CG | 32 (47.06) | 2 (6.25) | 0 (0.00) | 11 (34.38) | 9 (28.13) | 10 (31.25) |
| CHG | 46 (54.76) | 4 (8.70) | 1 (2.17) | 13 (28.26) | 16 (34.78) | 12 (26.09) |
| CHH | 863 (64.45) | 175 (20.28) | 21 (2.43) | 286 (33.14) | 252 (29.20) | 129 (14.95) |

Finally, we conducted a search for DMRs overlapping with 1,577 *B. cinerea* genes with known targeted functions that was carried out in the resequencing section above. Of these, 478 genes (30.3%) overlapped with one or more detected DMRs (See Table 1 and Table S8 for a comprehensive list of genes overlapping with DMRs).

## Discussion

### Culture-induced changes to MSAP profiles and virulence are simultaneous and reversible

In accordance with previous reports (Akamatsu *et al.*, 1999; Chu *et al.*, 2002; Hatta *et al.*, 2002; Castro *et al.*, 2003), virulence of *B. cinerea* cultures progressively decreased with culture age, but recovered after one passage of *in vivo* infection on *A. thaliana* (Butt *et al.*, 2006). Concurrent with these changes, MSAP profiles showed a similar progressive increase in deviation from the starter culture profiles consistent with previous reports of accumulative genetic/methylome change for other species, when similarly exposed to prolonged periods of culture (Rodríguez López *et al.*, 2010). This accumulation of variation as culture progressed showed a positive linear correlation with the observed changes in virulence. This accords with reported high levels of somaclonal variability arising during *in vitro* growth of phytopathogenic fungi, which seemingly depresses the level of virulence of the culture isolates (Dahmen *et al.*, 1983). Whilst the source of the increased variation during protracted culture could have a genetic or epigenetic cause (both mutation and methylation perturb MSAP profiles), reversal of virulence phenotype after a single passage of the cultured fungus on *A. thaliana* (T8P) strongly implies that a plastic, epigenetic mechanism is primarily responsible for driving the observed decline in virulence. This is most easily explained if culture alters the methylation status of the genome and this in turn perturbs the expression control of genes responsible for virulence. For this hypothesis to hold, a particularly strong link should exist between changes in the methylation status of genes known to be implicated in virulence and the disease symptoms evoked by culture. Establishing such links requires a more precise description of both methylome flux and mutation in culture.

### Sequence variants do not explain loss of virulence during culture

Whole genome resequencing of six DNA samples taken at two time points (one month and eight months) allowed us to monitor for mutational change during the extended period of *B.* cinerea culture. This work uncovered 198 non-synonymous mutations within genes. For genes with unknown function, it is difficult to implicate these changes as being involved in the loss of virulence observed during culture. We therefore screened specifically for the appearance of genetic variants among 1,577 *B. cinerea*. Of these, just eight genes (0.5%) included sequence variants that were not silent, conservative missense or synonymous mutations. Of the 1184 genes associated to virulence, just six (0.5%) included a sequence variant that could conceivably affect the virulence phenotype. All six were plant cell wall disassembly genes, CAZyme genes (Lombard *et al.*, 2014). It is tempting to speculate that some of the loss of virulence may have been casually linked to one or more of these loss-of-function mutants being favored *in vitro* such that virulence was progressively eroded but later restored by strong selection on the plant for functioning alleles. However, there is extensive redundancy among the 275 CAZyme genes present in the Botrytis genome (Blanco-Ullate *et al.,* 2014) and it is highly unlikely that the gradual loss of one or a small number of these genes would have such a pronounced effect on the disease phenotype. Equally, it is difficult to reconcile the stochastic and infrequent nature of mutation against the consistency in the timing and extent of virulence erosion/restoration seen across all replicates. Thus, we are inclined not to concur with the view of previous studies for other species that genetic mutation is the primary cause of the erosion of virulence during *in vitro* culture of *B. cinerea* (Jeon *et al.*, 2013).

### *In vitro* culture affects whole-genome methylation patterns

Our results suggest that changes in DNA methylation at a genome level are more likely than mutation to be causally linked to the observed loss of virulence. To establish a more direct association between methylome changes and gene activity we sequenced the methylomes of nine samples from three culture time points (T1, T8 and T8P). All genomes presented similarly low levels of DNA methylation (0.6%) as reported for other fungal species (Zemach *et al.*, 2010). DNA methylation was not evenly distributed across contigs but clustered following a mosaic pattern (Feng *et al.*, 2010), with higher levels of methylation in genomic regions with low gene abundance. Jeon *et al.*, (2015) noted similar methylation clustering patterns in pathogenic fungi around gene-poor regions rich in transposable elements rich and found these to dynamically follow fungal development. However, in our study clustering patterns were similar across time points suggesting that the methylation changes detected here are unlikely to be associated to any developmental progression that occurred during culture. However, global methylation levels varied with sequence context (CHH>CG>CHG) and time in culture. In brief, methylation on CGs and CHGs increased significantly between 1 (T1) and 8 (T8) months in culture. In both sequence contexts, following inoculation on *A. thaliana*, T8P cultures lost much of the methylation accumulated during culture and were no longer significantly different to the virgin inoculums. This pattern of methylation flux concords with our MSAP results and indicates that methylome variation accumulates in culture but is lost after contact with the host.

Previous genome-wide studies have shown that gene methylation in the monophyletic Ascomycota phylum varies greatly. For example, in *Neurospora crassa* DNA methylation is not found in gene bodies. However gene body methylation has been reported in other species such as, *Candida albicans*, *Uncinocarpus reesii* and *Magnaporthe oryza* (Zemach *et al.*, 2010; Su *et al.*, 2011; Jeon *et al.*, 2015) while promoter methylation has only been reported in *M. oryza* (Jeon *et al.*, 2015). Analysis of the effect of methylation distribution and density on gene expression has shown that gene body methylation has positive effects on gene expression (Zemach *et al.*, 2010; Jeon *et al.*, 2015) while promoter methylation has a negative effect (Jeon *et al.*, 2015). Our analysis of methylation distribution in *B. cinerea* genes showed an increase in methylation approximately 800bp upstream of the TSS followed by a sharp decrease at the TSS as shown by Jeon et al (2015) in *M. oryza* mycelia. Our results showed a second increase in methylation density after the TSS indicating that at least some *B. cinerea* genes contain gene body methylation.

Comparative analysis between culture time points showed that when all *B. cinerea* genes were analysed collectively there were no changes in DNA methylation with time in culture. However, while DNA methylation on house-keeping remained stable between culture time points, CAZyme genes showed an increase in promoter methylation with time in culture. Moreover, CAZyme genes showed a return to original methylation levels after a single passage on *A. thaliana* (i.e. T1=T8P<T8). Conversely, all *B. cinerea* genes analysed collectively showed an increase in methylation, similar to that observed at a whole genome scale (T1=T8<T8P). Global methylation levels on the CAZyme gene in promoter regions showed a negative correlation with virulence levels. This observed increase in promoter methylation on T8 samples seems to be due to an increase on the CHG and CG contexts. Conversely, global methylation levels within CAZyme gene bodies were positively correlated with virulence levels, largely due to changes in the CHH context. The CAZyme gene family encodes proteins that breakdown, biosynthesise and modify plant cell wall components and are highly expressed during plant invasion in *B. cinerea* (Lombard *et al.*, 2014; Blanco-Ulate *et al.*, 2014). This gene family, should therefore, be readily accessible to the transcriptional machinery in the virulent form of pathogenic fungi. Interestingly, our results show an association between virulence levels in T1, T8 and T8P samples and a methylation features in CAZyme genes that have positive effects on gene expression, i.e. high gene-body methylation (Zemach *et al.*, 2010; Jeon *et al.*, 2015) and low promoter methylation (Jeon *et al.*, 2015) in high virulence cultures (T1 and T8P) and the opposite in the low virulence culture (T8).

### Culture induced DMRs are reversible and mirror changes in virulence

Overall, we identified 2,822 candidate DMRs between T1, T8 and T8P for all mCs, with DMRs showing a decrease in methylation between T1 to T8 in all contexts (CG, CHG and CHH), followed by a recovery of DNA methylation after culture on *A. thaliana* (T8P). Initially, this seems to contradictobservations in global DNA methylation levels in which methylation increased following the trend (T1<T8<T8P). This might simply suggest that there is a marked difference between the behaviour of clustered and dispersed methylation. This assertion certainly concurs with similar differences noted between global ad local DNA methylation levels in previous studies on different organisms (Jung & Pfeifer, 2015; Jeon *et al.*, 2015). However, it should be noted that T8 cultures exhibited a larger number of outlier DMRs showing containing raised methylation levels, suggesting that not all predicted DMRs followed the same pattern of demethylation followed by re-methylation. Furthermore, regional changes in DNA methylation have been previously associated to the developmental potency of fungal cells (Jeon *et al.*, 2015). In their work, the authors showed how fungal totipotent cells (mycelia) possess higher global methylation levels while cells determined to host penetration (appresoria) contain a higher number of genes with methylated cytosines but lower global levels of DNA methylation. More pertinently, 68.3% of the culture-induced DNA methylation changes (i.e. Type 0, 2 and 1a, which accounts for 57.3% of the total detected DMRs) showed a recovery pattern after a single round of inoculation on *A. thaliana*. This suggests that a majority of the DNA methylation changes accumulated during *in vitro* culture are reset to their original state after a single passage on the host. Viewed in this context, 26.7% of the observed changes (Type 1b) were not induced by *in vitro* culture but by exposure to the host. This highlights the importance of epigenetic mechanisms in regulatinghost/pathogen interactions (Gómez-Díaz *et al.*, 2012; Weiberg *et al.*, 2013; Cheeseman & Weitzman, 2015), and the potential implication of dynamic DNA methylation as regulator of phenotypic plasticity in response to nutrient availability and interaction with the host (Mishra *et al.*, 2011).

Of the detected DMRs for all mCs 84.5% overlapped with one or more genes suggesting that the great majority of culture-induced DNA methylation flux occurs in genic regions. This overlap of individual DMRs with one than more gene is probably partially due to the small average size of intergenic regions (778 to 958bp) and genes (744 to 804 bp) in *B. cinerea* (Amselem *et al.*, 2011). Moreover, 98% of these overlapped, at least partially, with gene bodies. Taken collectively, this indicates that *B. cinerea* genes in general and gene bodies in particular are prone to environmentally-induced change to their methylation status. When DMRs were defined using each DNA methylation context independently (i.e. CG, CHG and CHH), the vast majority (89.1% of all DMRs and 89.9% of those overlapping with genes) occurred in the CHH context, implying that DMRs in this context are the reason for the observed changes in detected CAZyme gene body methylation associated with higher levels of virulence. Analysis of DMRs overlapping with genes with known function revealed that house-keeping genes did not overlap with any DMRs while genes associated with apoptosis and conidiation showed the higher percentage of overlapping DMRs (40.0 and 37.5% respectively). Remarkably, both biological processes have been previously shown to be affected by DNA methylation (Belden *et al.*, 2011; Hervouet *et al.*, 2013; Jeon *et al.*, 2015). More importantly 27.6% of all virulence genes analyzed overlapped with DMRs of which 77% showed recovery after a single passage on the host.

Our results imply that protracted culture of *B. cinerea* induces a hypermethylated, low pathogenic form adapted to the host’s absence or the abundance of nutrients in the culture media. It is tempting to speculate that the observed change in global and local levels of DNA methylation during *in vitro* culture could be part of a mechanism that confers plasticity to the *B. cinerea* genome to adapt to different environments. Butt *et al.,* (2006) proposed that *in vitro* culture induced loss of virulence could be the reflexion of an adaptive trait selected to promote energy efficiency, by turning off virulence genes in the absence of the host or in nutrient rich environments, and that such change could become maladaptive by restricting the pathogen to saprophytism (Butt *et al.*, 2006). Environmentally induced epigenetic adaptive changes have been predicted to potentially induce evolutionary traps leading to maladaptation (Consuegra del Olmo & Rodriguez Lopez, 2016). However, the complete reversibility of the reduced virulence phenotype suggests that this might not be the case here.

We propose that the next step towards a better understanding of the epigenetic regulation of virulence in fungi should be to confirm the causal links between the methylation changes described here and perturbations in virulence genes expression. This, together with the increasing availability of sequenced fungal genomes and methylomes, and our ability to decipher the associations between changes in DNA methylation and virulence, will stimulate the understanding of the mechanisms involved in fungal virulence control. Moreover, the analysis of DMRs could potentially be used to predict gene function in non-model fungal species or even to predict pathogenicity. More importantly, if the epigenetic regulation of the virulent/non-virulent transition proposed here applies broadly to other pathogenic fungi, our findings will open the door to a new type of non-lethal fungicide aimed at maintaining pathogenic fungi as saprotrophic that would reduce the appearance of resistant strains.

## Author contributions

J.B. did sequence data analysis and helped drafting the manuscript. A.S. cultured the *Botrytis* samples and performed the virulence analysis. A.A. undertook the estimations of *in planta* fungal develop using qPCR. M.W. and L.A.J.M. helped to conceive the study and drafting the manuscript. C.M.R.L. conceived the study, carried out all nucleic acid extractions, performed and analysed the MSAPs, performed and assisted analysing the native and bisulfite treated DNA sequencing and wrote the manuscript. All authors have read and approved the final manuscript.

## Supplementary Information

**Fig. S1** Schematic representation of experimental design

**Fig. S1.**
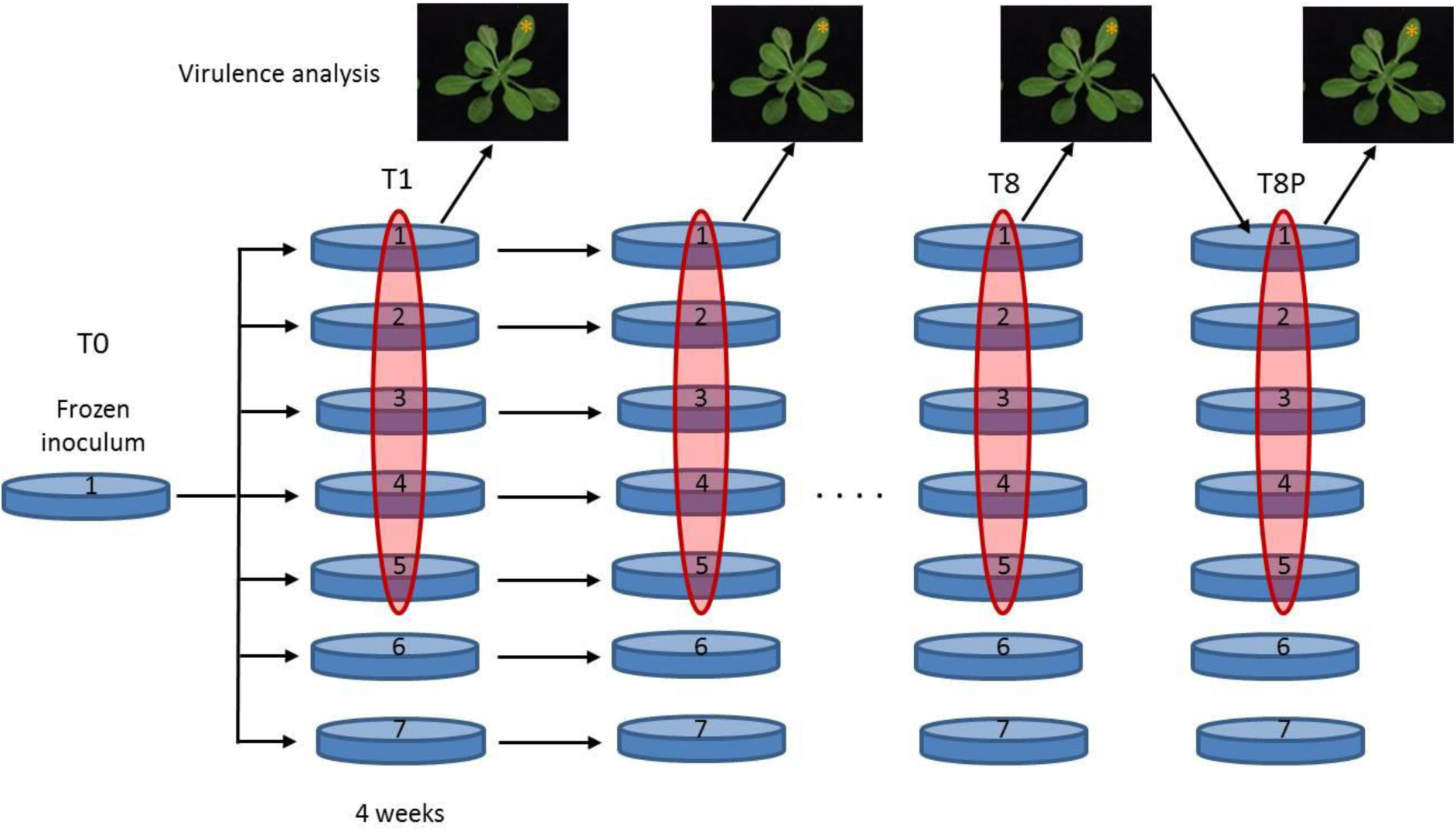
Seven replicated *Botrytis cinerea* cultures were initiated from a single frozen inoculum and cultured for 36 weeks. All replicates were subcultured every 4 weeks to fresh culture medium and mycelia were kept for DNA analysis (MSAPs, WGS, and BS-WGS). Conidia from five randomly selected replicates (highlighted in red) at each time point were collected and used for virulence analysis by inoculating five *Arabidopsis thaliana*. Inoculated *A. thaliana* leaves* at T1, T8 and T8P were collected 72 hours after inoculation for estimation of *in planta* fungal development by quantitative PCR. Conidia from were collected from *A. thaliana* tissue infected with T8 fungus and were re-cultured for a short time to generate conidia to test virulence recovery.

**Fig. S2** Correlation between pairwise molecular differentiation and changes in virulence observed during *in vitro culture* of *B. cinerea*.

**Fig. S2.**
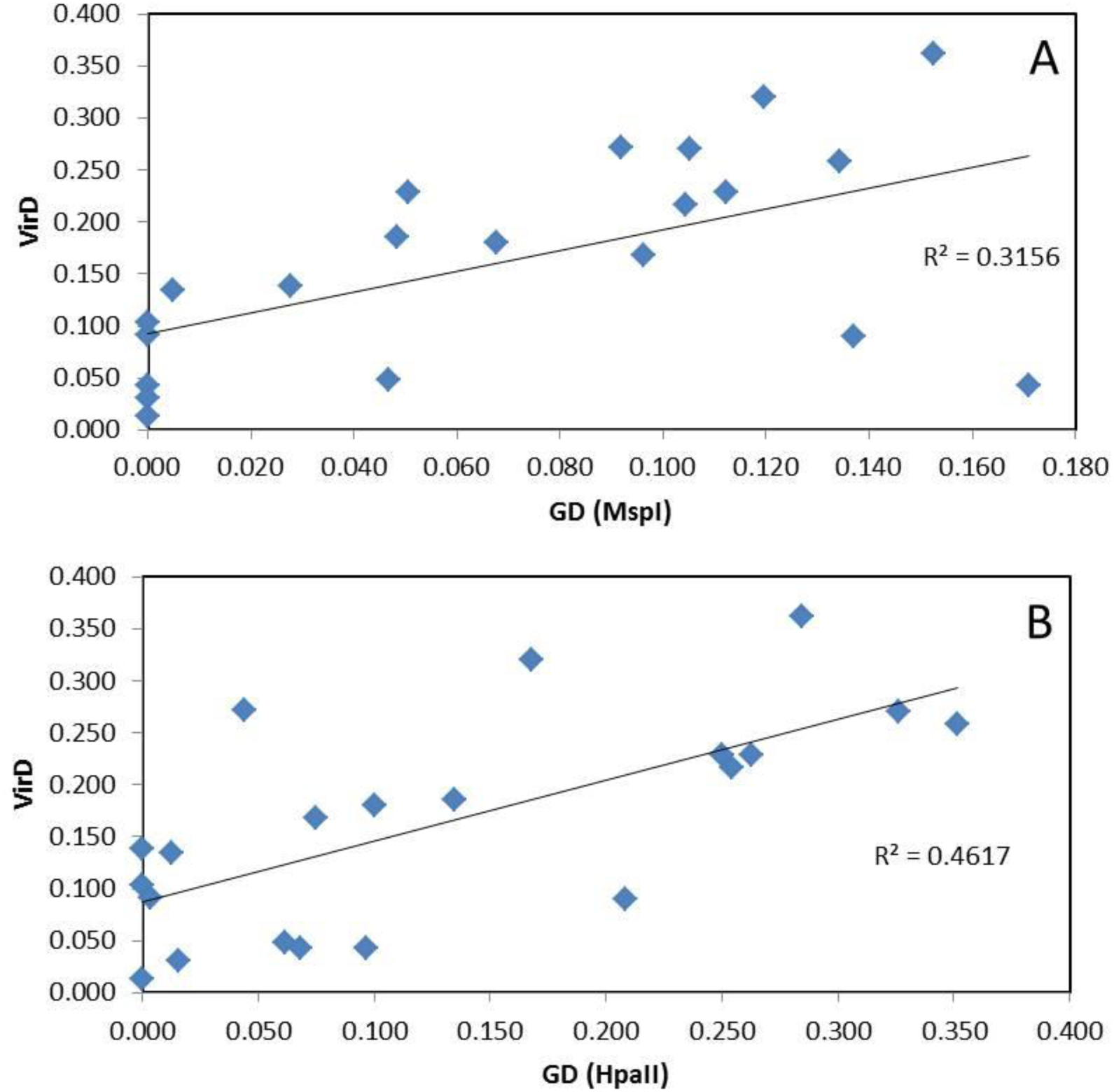
Correlations were calculated using mantle test analysis on pairwise PhiPT values **(GD)** (generated from MSAP data obtained using MspI **(a)** and HpaII **(b)**) and differences in virulence between culture time points **(VirD)**. Analysis using 1000 permutations showed significant correlations (**a**: P = 0.005; **b**: P = 0.002).

**Table S1** Sequences of oligonucleotide used for MSAP and qPCR.

**Table S1.**
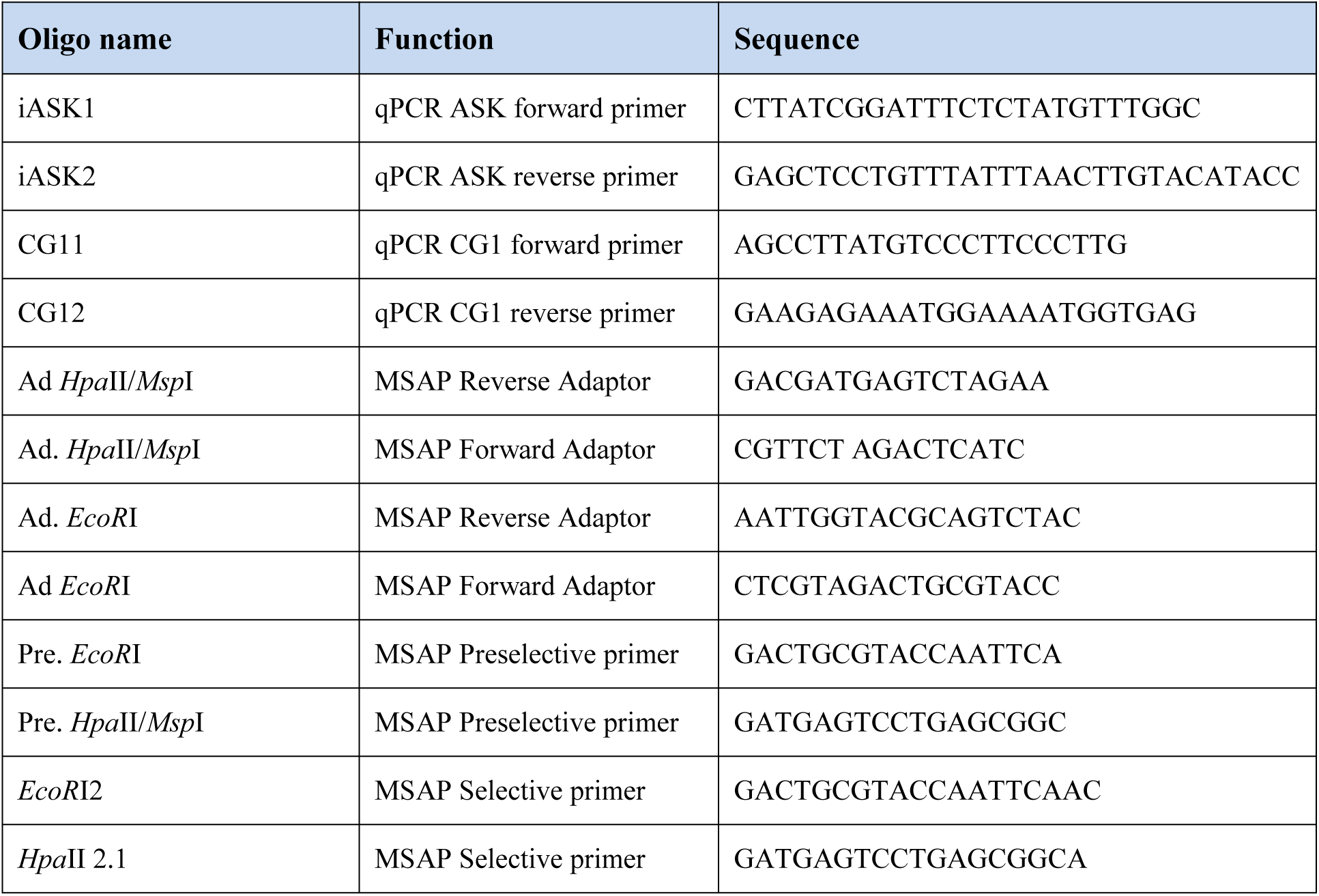
Click here to enter text.

**Table S2** B. cinerea genes presenting sequence variants between 1 (T1) and 8 (T8) month old cultures.

**Table S2** + indicates the functional group the gene carrying the sequence variant belongs to. Some genes present more than one sequence variant and can be present in more than one functional group. Variant categories were analysed using CooVar, bedtools and custom scripts.

**Table S3** Average coverage statistics for Whole Genome Bisulfite Sequenced samples.

**Table S3.** Average measurements were obtained from three replicates at each time point. Sequencing reads from all samples were mapped to *B. ciniera* B05.10 genome using Bismark/Bowtie2.

| Sample | Mean Coverage | Std dev | Cytosine coverage | Cytosine coverage (>4x) | mC (>1x) |
| --- | --- | --- | --- | --- | --- |
| T1 | 54.9939 | 20.3428 | 92.55% | 87.65% | 10.99% |
| T8 | 33.1991 | 13.868 | 91.50% | 81.07% | 9.14% |
| T8P | 45.0209 | 17.0144 | 93.42% | 86.67% | 10.63% |

**Table S4 Genome mapping statistics from 9 bisulfite converted Botrytis samples.**

**Table S4.** Sequencing reads from all samples were mapped to *B. ciniera* B05.10 genome using Bismark/Bowtie2

| Samples | Total bp | % Map. Effic | Total methylated C's |  |  | Total non-methylated C's |  |  | % methylation |  |  | %mC |
| --- | --- | --- | --- | --- | --- | --- | --- | --- | --- | --- | --- | --- |
|  |  |  | CpG | CHG | CHH | CpG | CHG | CHH | CpG | CHG | CHH |  |
| T1BS1 | 26382612 | 67.6 | 477254 | 438983 | 2499975 | 130600424 | 111869938 | 391807290 | 0.36 | 0.39 | 0.63 | 0.54 |
| T1BS2 | 34048060 | 62.9 | 526341 | 485477 | 2494454 | 155155501 | 134341879 | 485255169 | 0.34 | 0.36 | 0.51 | 0.45 |
| T1BS3 | 17347608 | 61.1 | 306505 | 279816 | 1575956 | 77226256 | 66054702 | 232502707 | 0.40 | 0.42 | 0.67 | 0.57 |
| AvT1 | 25926093.33 | 63.87 | 436700.00 | 401425.33 | 2190128.33 | 120994060.33 | 104088839.67 | 369855055.33 | 0.36 | 0.38 | 0.59 | 0.52 |
| T8BS1 | 18543427 | 62 | 394348 | 360641 | 1568745 | 83351098 | 71894819 | 257263175 | 0.47 | 0.50 | 0.61 | 0.56 |
| T8BS2 | 12613134 | 60.7 | 248968 | 229853 | 902624 | 55786373 | 47759672 | 166229028 | 0.44 | 0.48 | 0.54 | 0.51 |
| T8BS3 | 17117307 | 65.1 | 391446 | 360065 | 2142019 | 79537475 | 69135859 | 257486354 | 0.49 | 0.52 | 0.83 | 0.71 |
| AvT8 | 16091289.33 | 62.60 | 344920.67 | 316853.00 | 1537796.00 | 72891648.67 | 62930116.67 | 226992852.33 | 0.47 | 0.50 | 0.67 | 0.59 |
| T8PBS1 | 28336888 | 62.3 | 524875 | 487874 | 3561078 | 126943754 | 110309124 | 404488048 | 0.41 | 0.44 | 0.87 | 0.71 |
| T8PBS2 | 18471780 | 66.9 | 435528 | 403984 | 2932991 | 89352607 | 77332840 | 279892313 | 0.49 | 0.52 | 1.04 | 0.84 |
| T8PBS3 | 14694194 | 54.3 | 220458 | 201667 | 924438 | 57483914 | 49293008 | 173794940 | 0.38 | 0.41 | 0.53 | 0.48 |
| AvT8P | 20500954.00 | 61.17 | 393620.33 | 364508.33 | 2472835.67 | 91260091.67 | 78978324.00 | 286058433.67 | 0.43 | 0.46 | 0.86 | 0.68 |

**Table S5** *Botrytis cinerea CAZYme genes selected for the analysis of methylation density shown on Figure 5.*

**Table S5** (Modified from Blanco-Ulate et al 2014)

**Table S6** In vitro culture and plant induced and differentially Methylated Regions (DMRs) in Botrytis cinerea.

**Table S6** DMRs were determined from whole genome bisulfite sequencing data using 3 sample Kruskal-Wallis test on a sliding window analysis conducted through swDMR. Methylation levels (%mC) were calculated as proportion of methylation (i.e., number of methylated Cs divided by the total number of Cs) and were considered significantly different when the calculated FDR was lower than 0.01. Three replicates were used per time point (T0, T8 and T8P, 2, 8 month in culture and 8 months and inoculation onto *A. thaliana respectively*). Samples were grouped according to four patterns on DNA methylation behavior (2, T0=T8P > T8; 1a, T0>T8=T8P; 1b, T0=T8<T8P and 0, T0=T8P < T8). Coverage indicates the calculated average sequencing coverage for each DMR.

**Table S7** *Botrytis cinerea in vitro induced Differentially Methylated Regions overlapping with genes.*

**Table S7** DRMs overlapping with genes (i.e., regions presenting significantly different methylation levels between one sample and the other two samples (FDR<0.01)) were determined using a three sample Kruskal-Wallis test. Methylation levels were analyzed by sliding window analysis using swDMR. DMRs were determined for all context. DMRs were grouped according to the genic region they overlapped with (i.e, Promoter, promoter and Gene body, promoter, Gene body and 3’UTR, gene body and 3’UTR and gene body). Methylation pattern was considered as recovered when methylation at T0 was equal (not significantly different) to T8P and lower or higher than that at T8.

**Table S8** B. cinerea genes with known function overlapping with in vitro culture induced DMRs.

**Table S8** Colums mC, CG, CHG and CHH indicate what region of the gene overlaps with the DRM and in what Cytosine context the DMR is observed (mC= all Cytosines; CG, CHG and CHH). DRM type indicates what DNA methylation behavior the DMR showed when comparing all three samples (2, T1=T8P > T8; 1a, T1>T8=T8P; 1b, T1=T8<T8P and 0, T1=T8P < T8).

***in vitro culture* of *B. cinerea*.**

## References

Akamatsu H, Taga M, Kodama M, Johnson R, Otani H, Kohmoto K. 1999. Molecular karyotypes for Alternaria plant pathogens known to produce host-specific toxins. Current Genetics 35: 647–656.

Amselem J, Cuomo CA, van Kan JA, Viaud M, Benito EP, Couloux A, Coutinho PM, de Vries RP, Dyer PS, Fillinger S, et al. 2011. Genomic analysis of the necrotrophic fungal pathogens Sclerotinia sclerotiorum and Botrytis cinerea. PLoS Genetics 7: e1002230.

Armaleo D, Miao V. 1999. Symbiosis and DNA methylation in the Cladonia lichen fungus. Symbiosis 26: 143 – 163.

Belden WJ, Lewis ZA, Selker EU, Loros JJ, Dunlap JC. 2011. CHD1 remodels chromatin and influences transient DNA methylation at the clock gene frequency. PLoS Genetics 7: e1002166.

Bird A. 2007. Perceptions of epigenetics. Nature 447: 396–398.

Blanco-Ulate B, Morales-Cruz A, Amrine KC, Labavitch JM, Powell AL, Cantu D. 2014. Genome-wide transcriptional profiling of Botrytis cinerea genes targeting plant cell walls during infections of different hosts. Frontiers in plant science 5: 435.

Boyes DC, Zayed AM, Ascenzi R, McCaskill AJ, Hoffman NE, Davis KR, Görlach J. 2001. Growth stage-based phenotypic analysis of Arabidopsis: a model for high throughput functional genomics in plants. The Plant Cell 13: 1499–1510.

Boyko A, Kovalchuk I. 2011. Genetic and epigenetic effects of plant-pathogen interactions: an evolutionary perspective. Molecular Plant 4: 1014–1023.

Butt T, Wang C, Shah F, Hall R. 2006. Degeneration of entomogenous fungi. In: Eilenberg J, Hokkanen H, eds. An Ecological and Societal Approach to Biological Control. Dordrecht: Springer Netherlands, 213–226.

Caballero A, Quesada H, Rolán-Alvarez E. 2008. Impact of amplified fragment length polymorphism size homoplasy on the estimation of population genetic diversity and the detection of selective loci. Genetics 179: 539–554.

Castro M, Kramer K, Valdivia L, Ortiz S, Castillo A. 2003. A double-stranded RNA mycovirus confers hypovirulence-associated traits to Botrytis cinerea. FEMS Microbiology Letters 228: 87–91.

Cheeseman K, Weitzman JB. 2015. Host-parasite interactions: an intimate epigenetic relationship. Cellular Microbiology 17: 1121–1132.

Choquer M, Fournier E, Kunz C, Levis C, Pradier JM, Simon A, Viaud M. 2007. Botrytis cinerea virulence factors: new insights into a necrotrophic and polyphageous pathogen. FEMS Microbiology Letters 277: 1–10.

Chu YM, Jeon JJ, Yea SJ, Kim YH, Yun SH, Lee YW, Kim KH. 2002. Double-stranded RNA mycovirus from Fusarium graminearum. Applied and Environmental Microbiology 68: 2529–2534.

Consuegra del Olmo S, Rodriguez Lopez CM. Epigenetic-induced alterations in sex-ratios in response to climate change: an epigenetic trap? BioEssays. 38(9). DOI: 10.1002/bies.201600058

Dahmen H, Staub T, Schwinn FJ. 1983. Technique for Long-Term Preservation of Phytopathogenic Fungi in Liquid Nitrogen. Phytopathology 73: 241.

Excoffier L, Smouse PE, Quattro JM. 1992. Analysis of molecular variance inferred from metric distances among DNA haplotypes: application to human mitochondrial DNA restriction data. Genetics 131: 479–491.

Feng S, Cokus SJ, Zhang X, Chen PY, Bostick M, Goll MG, Hetzel J, Jain J, Strauss SH, Halpern ME, et al. 2010. Conservation and divergence of methylation patterning in plants and animals. Proceedings of the National Academy of Sciences of the United States of America 107: 8689–8694.

Fournier E, Gladieux P, Giraud T. 2013. The “Dr Jekyll and Mr Hyde fungus” : noble rot versus gray mold symptoms of Botrytis cinerea on grapes. Evolutionary applications 6: 960–969.

Gachon C, Saindrenan P. 2004. Real-time PCR monitoring of fungal development in Arabidopsis thaliana infected by Alternaria brassicicola and Botrytis cinerea. Plant Physiology and Biochemistry 42: 367–371.

Garrison E, Marth G. 2012. Haplotype-based variant detection from short-read sequencing.

Geyer KK, Rodríguez López CM, Chalmers IW, Munshi SE, Truscott M, Heald J, Wilkinson MJ, Hoffmann KF. 2011. Cytosine methylation regulates oviposition in the pathogenic blood fluke Schistosoma mansoni. Nature Communications 2: 424.

Gómez-Díaz E, Jordà M, Peinado MA, Rivero A. 2012. Epigenetics of host-pathogen interactions: the road ahead and the road behind. PLoS Pathogens 8: e1003007.

Gower JC. 1966. Some Distance Properties of Latent Root and Vector Methods Used in Multivariate Analysis. Biometrika.

Hatta R, Ito K, Hosaki Y, Tanaka T, Tanaka A, Yamamoto M, Akimitsu K, Tsuge T. 2002. A conditionally dispensable chromosome controls host-specific pathogenicity in the fungal plant pathogen Alternaria alternata. Genetics 161: 59–70.

He XJ, Chen T, Zhu JK. 2011. Regulation and function of DNA methylation in plants and animals. Cell Research 21: 442–465.

Hervouet E, Cheray M, Vallette FM, Cartron PF. 2013. DNA methylation and apoptosis resistance in cancer cells. Cells 2: 545–573.

Jeon J, Choi J, Lee GW, Dean RA, Lee YH. 2013. Experimental evolution reveals genome-wide spectrum and dynamics of mutations in the rice blast fungus, Magnaporthe oryzae. Plos One 8: e65416.

Jeon J, Choi J, Lee GW, Park SY, Huh A, Dean RA, Lee YH. 2015. Genome-wide profiling of DNA methylation provides insights into epigenetic regulation of fungal development in a plant pathogenic fungus, Magnaporthe oryzae. Scientific reports 5: 8567.

Johnson H, Broadhurst D, Goodacre R, Smith A. 2003. Metabolic fingerprinting of salt-stressed tomatoes. Phytochemistry 62: 919–928.

Johnson H, Lloyd A, Mur L, Smith A, Causton D. 2007. The application of MANOVA to analyse Arabidopsis thaliana metabolomic data from factorially designed experiments. Metabolomics?: Official journal of the Metabolomic Society 3: 517–530.

Jung M, Pfeifer GP. 2015. Aging and DNA methylation. BMC Biology 13: 7.

Jupe ER, Magill JM, Magill CW. 1986. Stage-specific DNA methylation in a fungal plant pathogen. Journal of Bacteriology 165: 420–423.

Kelly SA, Panhuis TM, Stoehr AM. 2012. Phenotypic plasticity: molecular mechanisms and adaptive significance. Comprehensive Physiology 2: 1417–1439.

Langmead B, Salzberg SL. 2012. Fast gapped-read alignment with Bowtie 2. Nature Methods 9: 357–359.

Li H, Handsaker B, Wysoker A, Fennell T, Ruan J, Homer N, Marth G, Abecasis G, Durbin R, 1000 Genome Project Data Processing Subgroup. 2009. The Sequence Alignment/Map format and SAMtools. Bioinformatics 25: 2078–2079.

Lloyd AJ, William Allwood J, Winder CL, Dunn WB, Heald JK, Cristescu SM, Sivakumaran A, Harren FJ, Mulema J, Denby K, et al. 2011. Metabolomic approaches reveal that cell wall modifications play a major role in ethylene-mediated resistance against Botrytis cinerea. The Plant Journal: for Cell and Molecular Biology 67: 852–868.

Lombard V, Golaconda Ramulu H, Drula E, Coutinho PM, Henrissat B. 2014. The carbohydrate-active enzymes database (CAZy) in 2013. Nucleic Acids Research 42: D490–D495.

Martin M. 2011. Cutadapt removes adapter sequences from high-throughput sequencing reads. EMBnet.journal 17: 10.

Martinez F, Dubos B, Fermaud M. 2005. The Role of Saprotrophy and Virulence in the Population Dynamics of Botrytis cinerea in Vineyards. Phytopathology 95: 692–700.

Michalakis Y, Excoffier L. 1996. A generic estimation of population subdivision using distances between alleles with special reference for microsatellite loci. Genetics 142: 1061–1064.

Mishra PK, Baum M, Carbon J. 2011. DNA methylation regulates phenotype-dependent transcriptional activity in Candida albicans. Proceedings of the National Academy of Sciences of the United States of America 108: 11965–11970.

Pathirana R, Cheah LH, Carimi F, Carra A. 2009. Low temperature stored in cryobank^®^ maintains pathogenicity in grapevine. cryoletters 30: 84.

Peakall R, Smouse PE. 2012. GenAlEx 6.5: genetic analysis in Excel. Population genetic software for teaching and research--an update. Bioinformatics 28: 2537–2539.

Quinlan AR, Hall IM. 2010. BEDTools: a flexible suite of utilities for comparing genomic features. Bioinformatics 26: 841–842.

Reyna-López GE, Simpson J, Ruiz-Herrera J. 1997. Differences in DNA methylation patterns are detectable during the dimorphic transition of fungi by amplification of restriction polymorphisms. Molecular & general genetics : MGG 253: 703–710.

Rodríguez López C, Morán P, Lago F, Espiñeira M, Beckmann M, Consuegra S. 2012. Detection and quantification of tissue of origin in salmon and veal products using methylation sensitive AFLPs. Food chemistry 131: 1493–1498.

Rodríguez López CM, Wetten AC, Wilkinson MJ. 2010. Progressive erosion of genetic and epigenetic variation in callus-derived cocoa (Theobroma cacao) plants. The New Phytologist 186: 856–868.

Rodríguez López CM, Wilkinson MJ. 2015. Epi-fingerprinting and epi-interventions for improved crop production and food quality. Frontiers in plant science 6: 397.

Su Z, Han L, Zhao Z. 2011. Conservation and divergence of DNA methylation in eukaryotes: new insights from single base-resolution DNA methylomes. Epigenetics 6: 134–140.

Thomma BP, Eggermont K, Tierens KF, Broekaert WF. 1999. Requirement of functional ethylene-insensitive 2 gene for efficient resistance of Arabidopsis to infection by Botrytis cinerea. Plant Physiology 121: 1093–1102.

Tricker PJ, Gibbings JG, Rodríguez López CM, Hadley P, Wilkinson MJ. 2012. Low relative humidity triggers RNA-directed de novo DNA methylation and suppression of genes controlling stomatal development. Journal of Experimental Botany 63: 3799–3813.

Vergara IA, Frech C, Chen N. 2012. CooVar: co-occurring variant analyzer. BMC Research Notes 5: 615.

Waddington C. 1942. Canalization of Development and the Inheritance of Acquired Characters. Nature 150: 563–565.

Weiberg A, Wang M, Lin FM, Zhao H, Zhang Z, Kaloshian I, Huang HD, Jin H. 2013. Fungal small RNAs suppress plant immunity by hijacking host RNA interference pathways. Science (New York) 342: 118–123.

Zemach A, McDaniel IE, Silva P, Zilberman D. 2010. Genome-wide evolutionary analysis of eukaryotic DNA methylation. Science (New York) 328: 916–919.

